# Genomic, transcriptomic, and metabolomic analysis of Traditional Chinese Medicine plant *Oldenlandia corymbosa* reveals the biosynthesis and mode of action of anti-cancer metabolites

**DOI:** 10.1101/2022.06.14.496066

**Authors:** Irene Julca, Daniela Mutwil-Anderwald, Vaishnervi Manoj, Zahra Khan, Soak Kuan Lai, Lay Kien Yang, Ing Tsyr Beh, Jerzy Dziekan, Yoon Pin Lim, Shen Kiat Lim, Yee Wen Low, Yuen In Lam, Yuguang Mu, Qiao Wen Tan, Przemyslaw Nuc, Le Min Choo, Gillian Khew, Loo Shining, Antony Kam, James P. Tam, Zbynek Bozdech, Maximilian Schmidt, Bjoern Usadel, Yoganathan s/o Kanagasundaram, Saleh Alseekh, Alisdair Fernie, Li Hoi Yeung, Marek Mutwil

## Abstract

Natural products from traditional medicinal plants are valuable candidates for clinical cancer therapy. Plants from the Oldenlandia-Hedyotis complex are popular ingredients of Traditional Chinese Medicine (TCM), however a major hurdle in the plant bioprospecting process of TCM plants is that the active metabolites, their biosynthetic pathways, and mode of action are often unknown. We show that *Oldenlandia corymbosa* extracts are active against breast cancer cell lines. To study the genes involved in the biosynthesis of active compounds in this medicinal plant, we assembled a high-quality genome. We show that the main active compound is ursolic acid and that abiotic stresses cause changes in anti-cancer activity, metabolite composition, and gene expression of plants. To reveal the mode of action of ursolic acid, we show that cancer cells undergo mitotic catastrophe, and we identify three high-confidence protein binding targets by Cellular Thermal Shift Assay (CETSA) and reverse docking.

## Introduction

Plants synthesize a vast array of secondary metabolites as part of their defense against pathogens or during stress conditions and are thus a natural resource for pharmaceuticals. This has been acknowledged already thousands of years ago by ancient medicinal systems such as traditional Chinese medicine (TCM) or Ayurveda. In the past years, massive efforts were made by researchers worldwide to mine this potential for new antibacterial or anti-tumor drugs ^1–4^. However, most studies are limited to screening the plants for activities and identifying candidates for the active metabolites without knowing the mode of action or the enzyme-coding genes. The major reason for this shortcoming is that the genomes of medicinal plants are usually not sequenced, and even if the genome sequence is known, the successful prediction of which enzyme-coding genes synthesize a given metabolite requires advanced computational methods applied to comprehensive datasets.

Plants already constitute a significant source of anti-cancer drugs approved by government health agencies such as the American FDA ^5,6^. The most well-known ones are taxol (paclitaxel), vinca alkaloids (vinblastine, vincristine), and camptothecin derivatives. The advantage of using natural products over chemotherapeutics is their effect on multiple signaling pathways and molecular targets. For example, they induce apoptosis, inhibit proliferation, and suppress metastasis, while causing few adverse effects ^3^. In contrast, classical chemotherapeutics are often single-targeted and therefore more prone to drug resistance ^7^. A recurring problem with medicinal herb extracts is that neither the active agents (and other molecular components of these mixtures) nor their modes of action are well-characterized. Additionally, the composition of the formulas is complex and not standardized, thus not fulfilling FDA guidelines for drug approval.

The Oldenlandia-Hedyotis complex of the family *Rubiaceae* comprises about 500 species found throughout the tropics. It is a highly polyphyletic group, and recent molecular phylogenetic studies suggest that it should be split into Hedyotis (robust, shrub-like species) and Oldenlandia (small herbs with paniculate or corymbose inflorescences) ^8^. *Oldenlandia corymbosa* is a commonly used herb in China and India for health benefits and wellbeing. Previous pharmacological studies reported that *O. corymbosa* has hepatoprotective ^9,10^, antimalarial ^11^, anti-inflammatory, and antioxidant properties ^12^. There are a few reports on anti-tumor activities of *O. corymbosa* against skin cancer, leukemia, and liver cancer cells ^13–15^. *O. corymbosa* is often the main ingredient in TCM herbal formulations such as the popular antitumor Peh-Hue-Juwa-Chi-Cao medicine together with *Oldenlandia diffusa* and other herbs ^12,16^.

Until these studies, *O. corymbosa* was long regarded as inferior when compared to the more widely studied *O. diffusa* (Li et al. 2013), which has been used for the treatment of inflammation-linked diseases, such as hepatitis, appendicitis, and urethritis in traditional Chinese medicine ^17^. *O. diffusa* also possesses strong anti-tumor activities against liver, lung, prostate, and ovarian cancers and is already used for clinical colon and breast cancer treatments in Taiwan ^18^. For example, *O. diffusa* ethanol extracts suppress the proliferation of colorectal cancer cells and induce cell apoptosis mediated by the suppression of the STAT3 pathway in mouse xenograft models and in human HT-29 cells ^19,20^. In breast cancer, *O. diffusa* is often used together with *Scutellaria barbata*, and treatment with these extracts results in inhibited proliferation and migration of three types of breast cancer cells in vitro as well as tumor growth in nude mice by apoptosis ^21^. *O. diffusa* extracts also exert antiproliferative and apoptotic effects on human breast cancer cells through ERα/Sp1-mediated p53 activation ^22^. Around 171 pharmacologically active compounds of *O. diffusa* have been reported ^17,23^. Thus, both *O. diffusa* and *O. corymbosa* have documented medicinal properties, but the identity, biosynthesis, and mode of action of the bioactive metabolites is still largely unknown.

Considering that natural products are increasingly popular in cancer therapy, we set out to identify the main anti-cancer metabolites and their biosynthetic pathways of the *Oldenlandia* genus. To this end, we collected 11 plants comprising *O. corymbosa*, *O. biflora,* and *O. tenelliflora*, and show that *O. corymbosa* had the most potent activity against several breast cancer cell lines. We provide a gene functional resource for *O. corymbosa*, by generating a high-quality genome sequence, which revealed a highly homozygous genome caused by autogamy. We used activity-guided fractionation to reveal that ursolic acid is the main metabolite responsible for anti-cancer activity, with a minor contribution of oleanolic acid, lutein, phytol, and pheophorbide L. The gene expression and metabolomic analysis of *O. corymbosa* organs showed that the biosynthesis of these metabolites can be influenced by abiotic stress. To study the mechanism of the anti-cancer activity of ursolic acid we show that it causes a mitotic catastrophe in the breast cancer cell lines. Finally, we identified three high-confidence human protein targets of ursolic acid by a combination of Cellular Thermal Shift Assay (CETSA) and reverse docking.

## Results

### Collection of *Oldenlandia* species and establishment of *O. corymbosa* in the laboratory

To study and compare the anti-tumor activities of different *Oldenlandia* species, we collected 10 field-grown plants from the Bedok area in Singapore and one plant from a private residential area (Table S1). The plants were transferred to the lab, grown on a typical soil substrate (7:2:1, soil:vermiculite:perlite), and propagated by selfing or clonal propagation. Then we classified the plants by morphological characteristics ^24^. *O. corymbosa* (M1, M2, M5, M11-13, M15) possesses lanceolate or elliptic leaves, the stems are branching near the base, and the inflorescences have 2-5 flowered corymbs or umbels (Figure 1A). *O. biflora* (M6, M7) leaves are elliptic-ovate, the flowers are cymose, 2−many flowered and situated terminal or in the axils of uppermost leaves, and the seed capsules are oblate (Figure 1B). *O. tenelliflora* (M9, M14) is characterized by having sessile, solitary flowers and laminar linear or linear-lanceolate leaves (Figure 1C).

**Figure 1.**
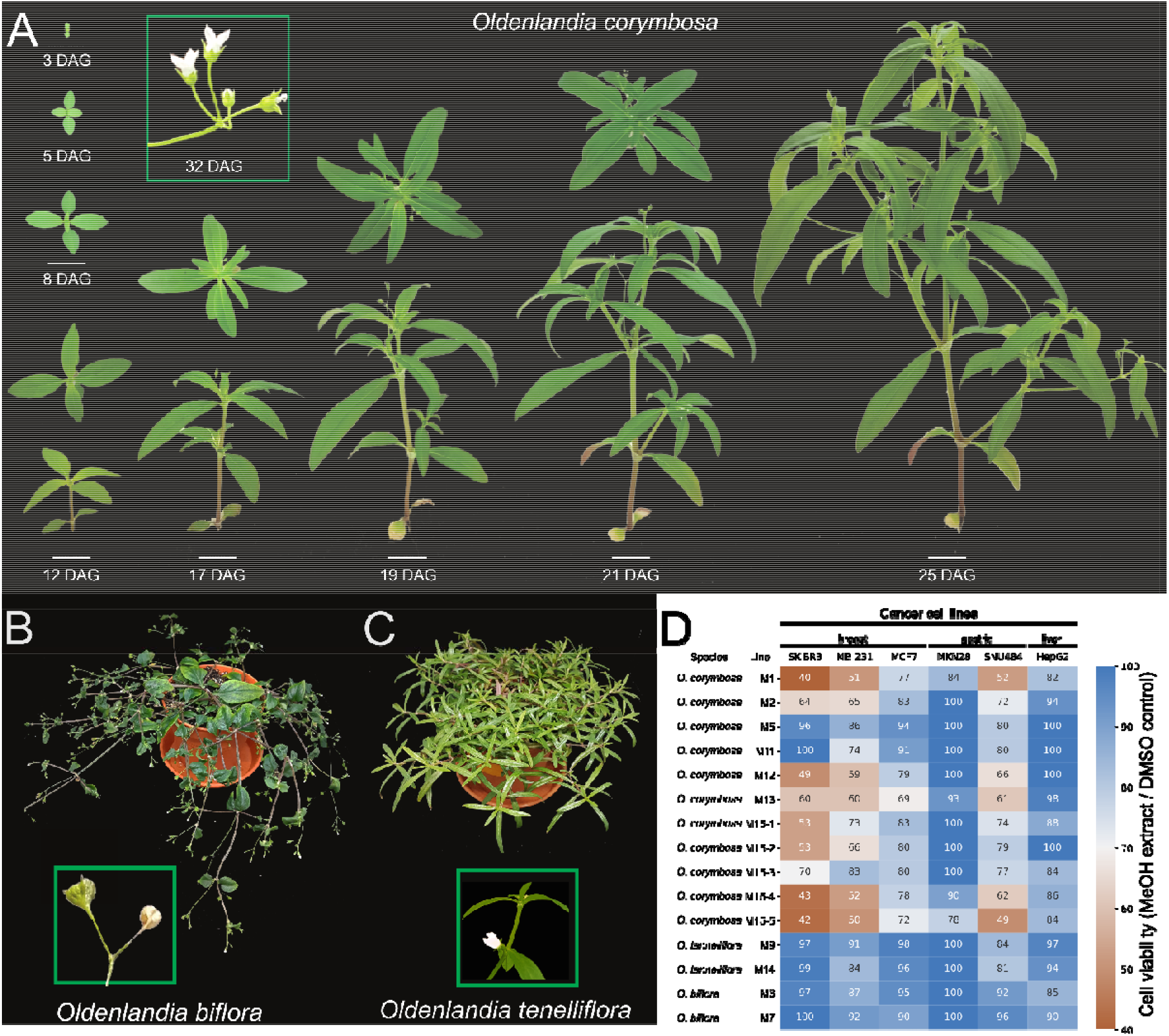
Identification and anti-tumor activities of Oldenlandia species. A) Growth of *O. corymbosa* over 25 days after germination (DAG). The images are taken from the top (DAG 3-8) and from the side (DAG 12-25). B) Morphological characteristics of *Oldenlandia biflora*. The bifurcating flowers are shown in the green box. C) *O. tenelliflora*, with the flower (green box). D) MTS assays showing the metabolic activity (proxy of viability) of six cancer cell lines (columns) and the various lines of *O. corymbosa* (M1, M2, M5, M11, M12, M13, F1 plants of M15), *O. tenelliflora* (M9, M14) and *O. biflora* (M6, M7). Values were capped at 100.

To select the species with anti-tumor activity, we tested methanol leaf extracts of the 11 plants against six cancer cell lines: breast cancer (SK-BR3, MB-231, MCF7), gastric cancer (MKN28, SNU484), and liver cancer (HepG2). To measure the cytotoxic activity of the methanol extracts we used the MTS cell viability assay which measures the metabolic activity of cancer cells, where lower metabolic activity is indicative of cell death ^25^. Nearly all leaf extracts from *O. corymbosa* plants showed strong activity against SKBR3 (∼40% cell viability, Figure 1D, Table S2), a somewhat lower activity (∼50%) against MB231 and SNU484, whereas MKN28, HepG2, and MCF7 were not or only modestly affected. The anti-tumor activity of *O. corymbosa* persisted after selfing the plant (Figure 1D, plants M15-1 to M15-5). Conversely, the extracts from *O. tenelliflora* and *O. biflora* showed modest or no activity against all six cancer lines (Figure 1D).

In addition to the anti-cancer activity, *O. corymbosa* grew rapidly, with the first flowers emerging ∼19 days after germination. The plant was readily self-fertilizing, able to be clonally propagated, grew well in soil, liquid, and solid media as per Arabidopsis protocols (Figure S1). The seeds were still viable after 3 years of storage (45% humidity, 23L). Thus, in addition to the anti-cancer activity, *O. corymbosa* shows all characteristics of a good model plant for the Oldenlandia genus.

To compare *O. corymbosa* to other medicinal plants regarding their activity against breast cancer, we performed 86 methanol extractions of 61 plants grown in Nanyang Technological University Herb Garden against MB231, SKBR3, and MCF7. Some of the extracts showed no activity (*Polygonum chinense*, *Rhoeo discolor*), or indiscriminately strong activity against all three cancer cell lines (Piper betle, Citrus hybrid, *Vitex rotundifolia*, *Wedelia chinensis*, *Clerodendratus spicatus*) (Figure S2). In contrast, *O. corymbosa* M15 plants showed more selective activity against SK-BR3 (Figure S2), suggesting that the plant extracts may target specific characteristics of this cancer line. Thus, we selected *O. corymbosa* and SKBR3 as our model plant and cancer cell line, respectively.

### The highly homozygous genome of *Oldenlandia corymbosa*

To enable molecular studies of *O. corymbosa* and to identify the biosynthetic pathways of the anti-tumor metabolites, we produced 79Gbp of long-read sequencing data from Oxford Nanopore Technologies (ONT) and 68Gbp of short reads from Illumina. The estimated genome size inferred from the *k*-mer frequency was 283.8Mbp (Figure S3a,b). The final assembly has 14 contigs (N50=31.05Mbp, Figure 2A), with a size of 274.6Mbp (Table 1), which represents 97% of the estimated genome size. The *k*-mer analysis also reveals a high completeness of the genome (99.8%, Figure S3c) with a consensus quality value (QV) of 38.5. The gene completeness of the assembly as determined by BUSCO (Benchmarking Universal Single-Copy Ortholog), showing 95.7% completeness (Table 1). This indicates high completeness comparable to other recently sequenced genomes ^26,27^. In addition, we assembled the plastid genome of *O. corymbosa*, which comprised 152,414 base pairs (bp), in agreement with previously reported *Oldenlandia* and *Hedyotis* plastid sequences, which range from 152,327 to 154,560 bp ^28,29^ (Table S3).

**Figure 2.**
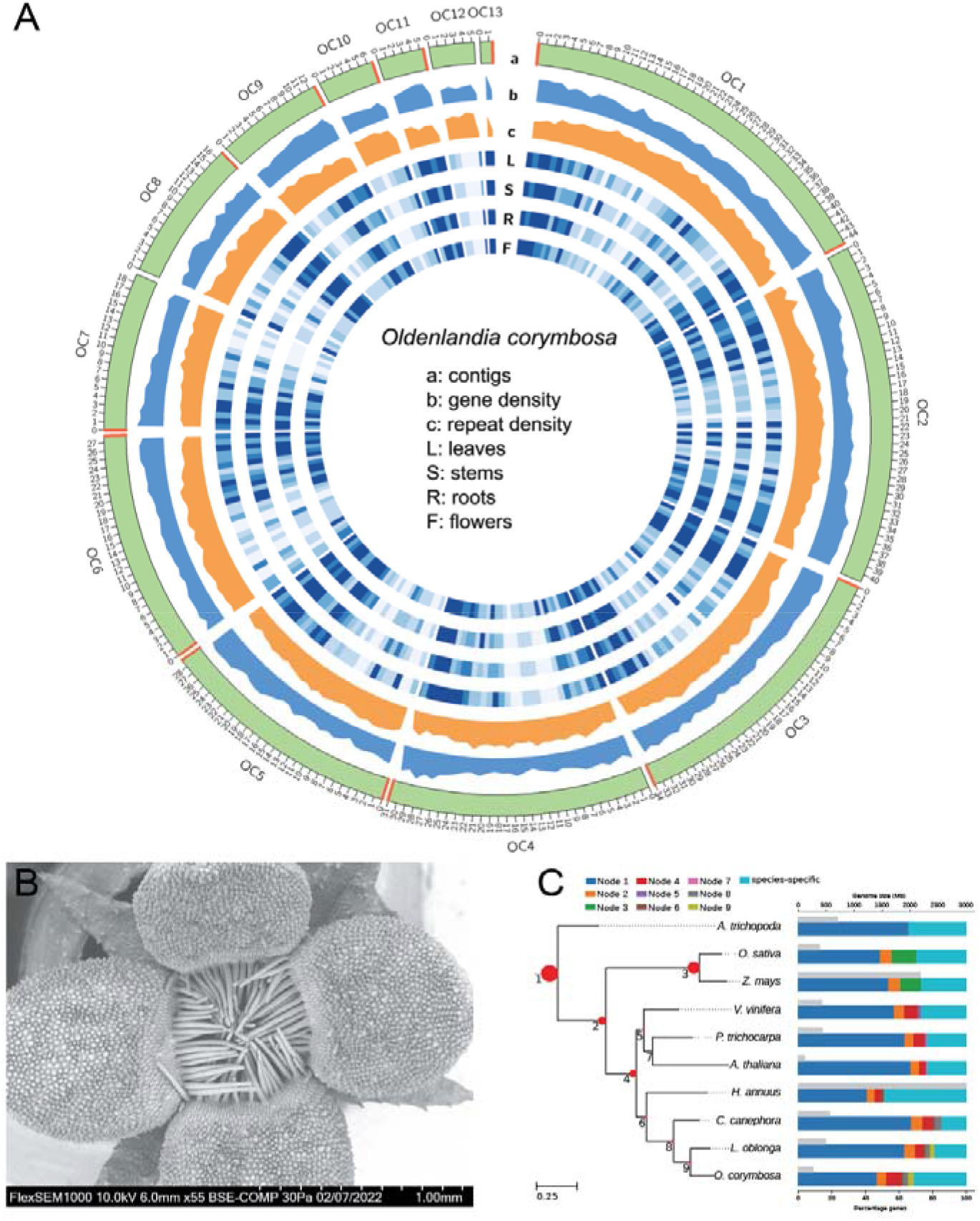
Genome characteristics of *Oldenlandia corymbosa*. A) Circos plot of the 13 largest contigs (green graph), gene density (blue graph), and repeat density (orange graph). The red regions in the contigs show the putative telomeres. The heatmaps in blue, in order, show the expression of genes in leaves, stems, roots, and flowers. B) Scanning electron microscopy (SEM) images of the opening of the flower. The scale bar (500µm) is indicated by white ticks. C) Species tree of *O. corymbosa* and other nine plant species. All bootstrap values in the graph are maximal (100). The numbers in the nodes indicate the node number (eg. 1: Node 1). The red circles in the tree show the percentage of orthogroups found in each node. Colour bars on the right show the percentage of genes per species that are present in each node. Grey bars on the right show the genome size (Mbp).

**Table 1.** Genome characteristics of *Oldenlandia corymbosa*

|  |  |
| --- | --- |
| GENOME SIZE | 274,568,628 |
| CONTIGS | 14 |
| CONTIGS > 1000kb | 14 |
| CONTIGS > 100kb | 13 |
| Largest scaffold | 44,422,723 |
| N50 | 31,047,018 |
| L50 | 4 |
| GC content | 0.36 |
| Number of N | 0 |
| BUSCO C | 95.70% |
| BUSCO S | 93.70% |
| BUSCO D | 2.00% |
| BUSCO F | 1.50% |
| BUSCO M | 2.80% |
| Number genes | 38,558 |
| Number of SNPs | 3,264 |
| Heterozygosity | 0.0012% |

To identify fully assembled chromosomes, we searched telomere repeats (5′- CCCTAAA-3′) ^30^ at both ends of the contigs and observed that contigs OC1, OC3, OC5, OC6 likely represent full chromosomes, as the telomeric sequences were observed on both ends (Figure 2A, telomere sequences indicated by red lines). Conversely, most of the other contigs contained at least one terminal telomere sequence. Interestingly, we also observed telomere-like sequences within the contigs (OC1, OC2, OC12). These internal sequences are called interstitial telomeric sequences (ITSs) and are thought to represent ancient chromosome fusions or remnants of double-stranded break repair events ^31,32^, and are often associated with genome instabilities ^33^. Altogether, these results show that our genome assembly has high continuity and completeness.

Surprisingly for a weed plant that was only selfed for one generation, the genome shows highly homozygosity levels as predicted by the k-mer analysis (99.9%, Figure S3a,b). Moreover, we only detected 3,264 heterozygous SNPs (Table 1, Figure S3b,c). To investigate this further, we performed a Scanning Electron Microscopy (SEM) imaging of mature flowers and observed that the entrance to the flower is blocked by a ring of pubescent hairs (Figure 2B). This characteristic may promote autogamy, which is reflected in the homozygosity observed in the genome.

The combined size of all repeat sequences identified in the genome of *O. corymbosa* is 87.9 Mbp (32% of the total genome length, Figure 2, Table S4). These repeats mainly consisted of retroelements (40 Mb, 46%) and unclassified interspersed repeats (32Mb, 37%). The total number of genes annotated is 38,558 based on de novo prediction, homology search, and mRNA-seq assisted prediction (Table 1). When we compare the repeat and gene density, we observe a general trend of gene density increasing and repeat density decreasing in some regions of the complete chromosomes (Figure 2b,c), which is typical for most flowering genomes ^34^. Additionally, we predicted functions for 26,221 genes (68%), including GO terms and MapMan bins to 18,423, and 13,231 protein-coding genes, respectively (Figure S3d). Finally, for the plastid genome, we annotated 131 genes, of which 86 are protein-coding genes, 37 are transfer RNAs, and eight are ribosomal RNAs (Figure S4a).

To study the evolutionary history of *O. corymbosa*, we compare its nuclear genome with other nine plant species and its plastid genome with other four Oldenlandia-Hedyotis species. The number of nuclear genes identified in *O. corymbosa* is higher than other Rubiaceae species but comparable to other plants (Table S5) and the number of plastid genes is comparable to other *Oldenlandia* species (Table S3). Also, for *O. corymbosa*, 26,599 nuclear genes (68%) have homologs in the other plant species, while for the other species this percentage ranges from 51% to 85% (Figure 2C). We reconstructed the evolutionary relationship of these 10 species using a concatenated approach with 807 single-copy orthologs, which resulted in a highly supported topology (Figure 2C) congruent with previous studies ^27^. Additionally, we reconstructed a phylogeny using the plastid genomes of Oldenlandia-Hedyotis species and *Coffea canephora* as an outgroup, which shows that our sample is sister to the other *O. corymbosa* included, strengthening the identity of our plant species (Figure S4b). The synteny analysis between *O. corymbosa* and *Leptodermis oblonga* indicates that O. corymbosa did not experience a whole-genome duplication event (Figure S5a,b). Similar results were reported for other Rubiaceae species ^27,35^.

### Identification of metabolites conferring the anti-cancer activity

Leaf extracts from *O. corymbosa* plants showed strong activity against the breast cancer line SKBR3 (Figure 1D, Figure S2). While metabolites are typically responsible for bioactivities of plants, also cyclotides, which are heat-stable macrocyclic peptides can be responsible for a wide range of activities. Cyclotides are likely important for plant defense and show activities that are anti-HIV ^36^, anti-cancer ^37^, antimicrobial ^38^, hemolysis ^39^, uterotensin ^40^, insecticidal ^41^, and nematicidal ^42^. Since cyclotides were found in *O. diffusa* ^37^ and *O. biflora* ^43^ we investigated the presence of these peptides of molecular weight between 2000 and 6000 Dalton (Da) in *O. corymbosa*. We observed a peak corresponding to a 3507 Da compound (Figure 3A, Figure S6, peak at 1753.52 m/z), which likely represents a cyclotide. To investigate whether the putative cyclotide is responsible for the anti-cancer activity, we compared the activity of the purified 3507 Da cyclotide to the 3507 Da-depleted fractions. MTT assays revealed that the cyclotide did not show any anti-cancer activity and that the cyclotide-depleted fraction exhibited activity (Figure 3B, Table S6), indicating that another compound is responsible for the activity.

**Figure 3.**
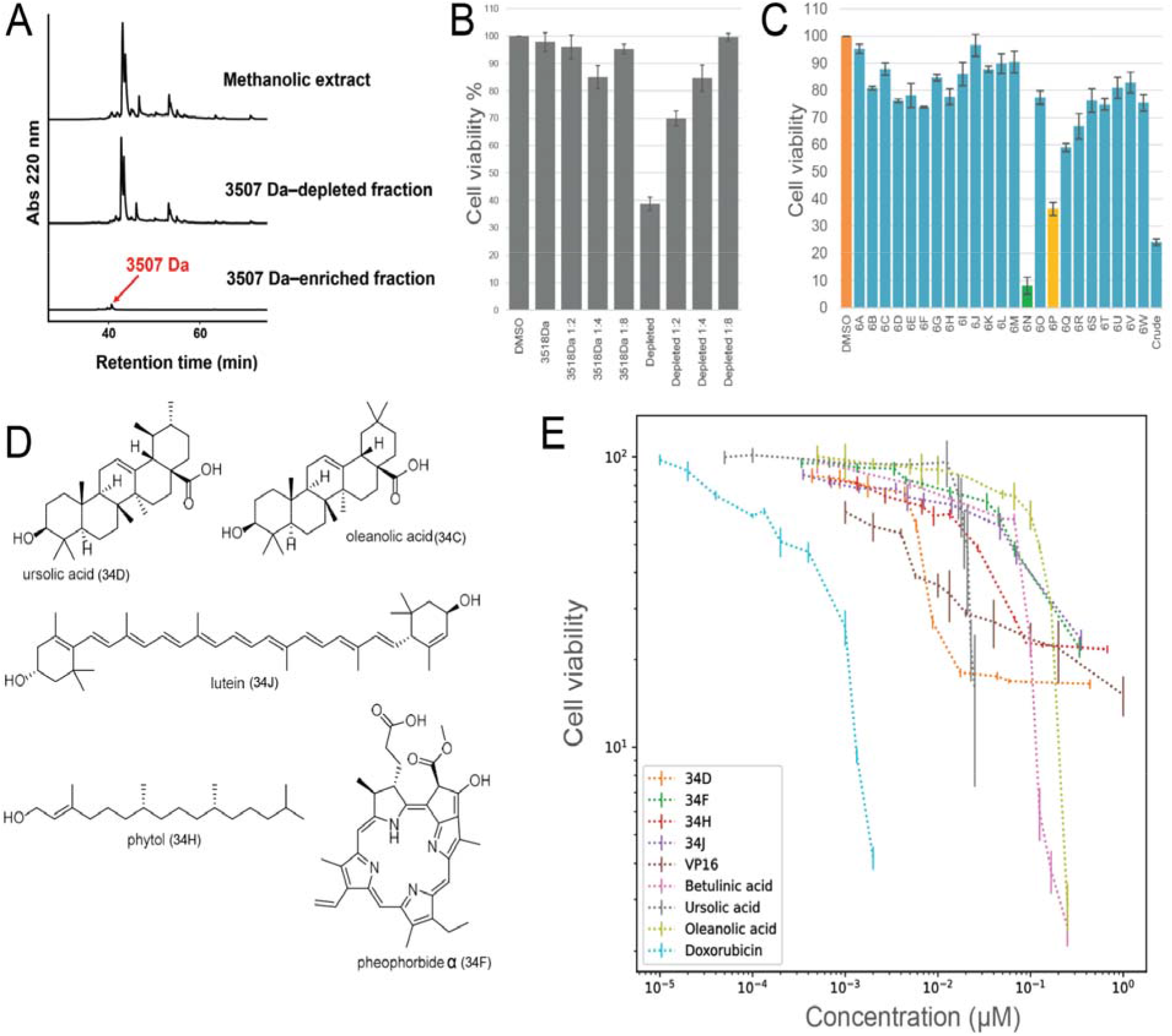
Identification of compounds of *O. corymbosa* with anti-tumor activities against SKBR3 cells. A) High-Performance Liquid Chromatograms of the methanolic *O. corymbosa* extract, and its 3507 Da-depleted and 3507 Da-enriched fractions at 220 nm. The red line corresponds to the peak containing the 3507 Da peptide. B) Cell viability assay of isolated peak and the depleted fraction. C) Cell viability assay of 23 fractions and crude control. D) Structure of the identified compounds. E) Dilution series for each fraction to determine IC50 values. Stock concentrations were 2 μg/μl for all fractions, i.e., 34D (4.38 mM), 34F (3.37 mM), 34H (6.745 mM), and 34J (3.5 mM). Stock concentrations for the commercially available compounds were ursolic acid (0.5 mM), oleanolic acid (OA, 5 mM), betulinic acid (BA, 5 mM), alkaloid etoposide (VP16, 10 mM), and anthracycline doxorubicin (Doxorubicin, 0.1 mM).

To identify the active compounds in *O. corymbosa* responsible for the inhibition of SKBR3 proliferation, we performed a comprehensive activity-guided fractionation and subjected 23 fractions (6A to 6W) together with the crude extract control to MTT assay analyses (Figure 3C). Two fractions (6N and 6P) showed strong activity against SKBR3 cells in addition to the crude extract control (Figure 3C, Table S7). High-resolution MS analysis of the fractions identified ursolic and oleanolic acid (fraction 6N), Pheophorbide L, phytol, and lutein (fraction 6P, Figure 3D) as putative active compounds. To pinpoint the active compounds, we performed a scale-up fractionation and confirmed the structures of the purified compounds by ^1^H NMR (Figure S7-8). The purified compounds (fractions 34D, F, H, J, see Table 2) were tested for the growth-inhibitory efficacy against SKBR3. To calculate the efficacy, we performed serial dilution to obtain the IC_50_ values (Figure 3E, Table S8), which indicate how much of a compound is needed to decrease the cell viability (captured by MTT assay) to 50%. We additionally included commercially available ursolic acid and its structural analogs oleanolic acid, and betulinic acid. To compare the activities of these compounds to common chemotherapy drugs we also included the alkaloid etoposide (VP16) ^44^ and the anthracycline doxorubicin ^45^.

**Table 2.** The computed IC50 range for the active fractions of *O. corymbosa* fo SKBR3 cancer cells and commercial compounds.

| Fraction/Standard | Purified amount (mg) | Best Fit IC<sub>50</sub> (µM) | Min (95% CI) | Max (95% CI) |
| --- | --- | --- | --- | --- |
| Ursolic acid (34D) | 0.6 | 6.681 | 5.225 | 8.536 |
| Pheophorbide **4** (34F) | 0.9 | 74.36 | 63.33 | 88.62 |
| Phytol (34H) | 2.6 | 20.33 | 16.92 | 24.49 |
| Lutein (34J) | 0.6 | 57.65 | 44.24 | 78.11 |
| Ursolic acid (Sigma) | - | 12.5 | 12.32 | 12.69 |
| Oleanolic acid (Sigma) | - | 109.7 | 96.15 | 123.8 |
| Betulinic acid (Sigma) | - | 66.65 | 59.7 | 73.34 |
| VP16 | - | 3.38 | 2.441 | 4.438 |
| Doxorubicin | - | 0.222 | 0.1875 | 0.263 |

The IC_50_ value of fraction 34D (6.7 µM) was comparable to the IC_50_ value of commercial ursolic acid (12.5 µM), but not to oleanolic (109.7 µM) or betulinic acid (66.65 µM), indicating that relatively minor modifications to a compound can have a large influence on the bioactivity. Phytol (20.33 µM), pheophorbide L (74.36 µM), and lutein (57.65 µM) showed lower activities than ursolic acid. Conversely, VP16 (3.38 µM) and anthracycline doxorubicin (0.22 µM) showed the most potent activities against SKBR3 cells. Taken together, the most active compounds in *O. corymbosa* are ursolic acid (6.8 µM) and phytol (20.33 µM).

### Abiotic stress modulates the biosynthesis of anti-tumor metabolites

The content of active compounds in medicinal plants can vary greatly depending on growth conditions and organs ^46,47^. To investigate how abiotic stress can influence the production of anti-tumor metabolites of *O. corymbosa*, we carried out a systematic approach involving eight abiotic stresses. We grew plants in soil under normal conditions (200 μmol·m^-2^·s^-1^, 28°C, 12h light) until DAG 23 (±2 days, until the 4^th^ node was developed, Figure S9A) and then transferred them to eight different stress conditions for 7 or 9 days. The 7-day long treatments included a short day (8h light/16h dark), long day (20h light/4h dark), cold (8°C and 15°C), and heat (40°C). The 9-day long treatments were drought (no watering) and two high light conditions (1000 and 2000 μmol·m^-2^·s^-1^).

The different stresses had impacts on overall plant morphology. The high light stresses increased the pigmentation of the plants with a stronger accumulation of the purple pigment at 2000 μmol·m^-2^·s^-1^, particularly in stems (Figure 4A). High light also led to thick and yellow leaves, more flowers, and a larger root system (Figure S9B). Cold treatment caused growth arrest, purple pigmentation of the stems at 15°C but not at 8°C, and rolled-in leaves (8°C, Figure S9C). 40°C heat treatment led to slow, more upright growth, few flowers, and yellow leaf color (chlorosis) of young tissues (Figure 4A). Short days resulted in light green, thin leaves (Figure 4A, 8L/16D) whereas plants grown in long days accumulated more yellow and purple pigments in leaves and stems and showed stunted growth. Finally, plants exposed to drought displayed a decrease in purple pigmentation, growth arrest, few flowers, and wrinkled leaves (Figure 4A). These results clearly show that the applied stresses have strong effects on the metabolism and growth of *O. corymbosa,* which should be reflected in metabolite changes.

**Figure 4.**
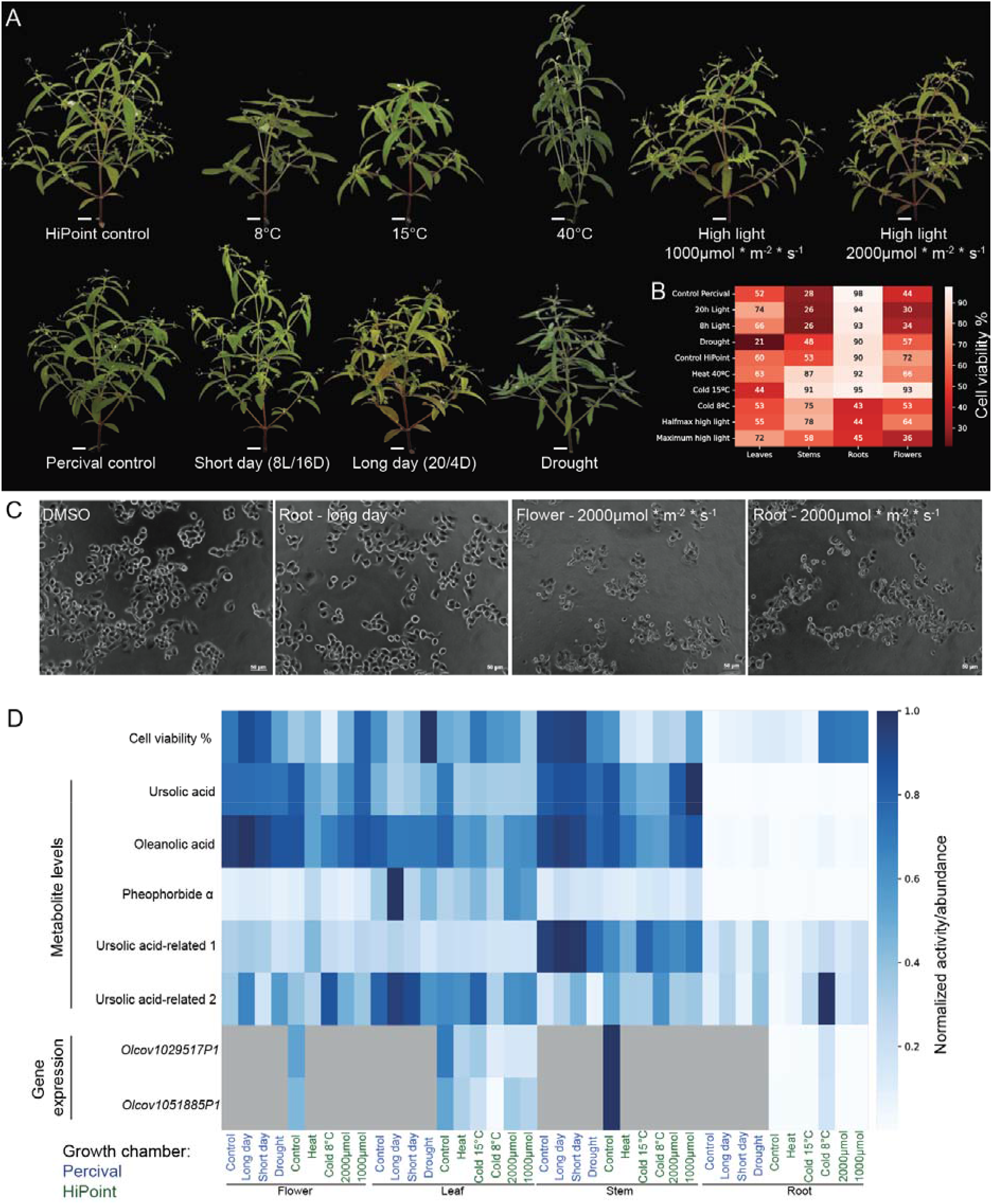
Abiotic stress treatment of *O. corymbosa*. A) Phenotypes of *O. corymbosa* plants grown for 7 days under the different abiotic stresses. B) MTT assay activities against SKBR3 cancer cell lines. The rows and columns represent abiotic stresses and organs, respectively. C) Phase-contrast images of SKBR3 cells untreated (DMSO control) and treated with metabolite extracts from roots from plants grown under long-day (20L/4D), high light flowers, and high light roots. D) Normalized activity and abundances across the different stress treatments. The rows indicate cell viability, the levels of metabolites, and gene expression, while the columns represent the different organs and abiotic stresses. The values are scaled to range from 0 (white cell color: no activity, absent metabolite, absent gene expression) to 1 (dark blue cell color: maximal activity, metabolite level, gene expression across all measurements). Green and blue letter colors represent samples grown in the Percival or HiPoint growth chamber, respectively.

To investigate the effects of the abiotic stresses on the anti-tumor potential, we performed MTT assays on root, leaf, stem, and flower extracts of the stressed plants (Table S9). We observed that leaves, stems, and flowers showed anti-tumor activity in most conditions (Figure 4B, 20-80% viability). Leave extracts of control plants showed ∼50% inhibition of SK-BR3 cells in line with our previous results (Figure 1D). Stem and flower extracts of the controls showed an inhibition range between 30 and 70% whereas root extracts did not show any activity for control conditions (>90% viability). Reduced activity of stem extracts was seen after cold and heat treatments (8°C, 15°C, and 40°C), and in high light (in four out of six biological replicates, Figure S10, Table S9). Drought increased activity (21%) of the leaf extract which could be due to water loss as suggested by the curling of leaves. In roots, after growth at 8°C, all biological replicates gained strong activity (43% viability). In the high light-treated plants half of the biological replicates gained anti-tumor activity in the roots (44-45%). To verify the viability status of the SKBR3 cells, we investigated their morphology by microscopy (Figure 4C). While cells treated with the DMSO control and the inactive root extract showed no evidence of cell death, cells treated with the root extracts from high light plants showed cell death (Figure 4C), similarly to cells treated by the usually active flower extract. Thus, these results suggest that the different abiotic stresses can modulate the metabolic composition of plants and anti-cancer activity.

### Metabolomic and gene expression analyses of biosynthetic pathways of anti-tumor compounds

We observed that abiotic stress modulated the anti-tumor activity of different organs, which could be a consequence of the altered activity of the metabolic pathways. To better understand this phenomenon, we quantified the relative abundance of ursolic acid, oleanolic acid, and pheophorbide L in the same methanol extractions that were used for MTT assays (Table S10). Lutein and phytol could not be quantified in these methanol extracts due to their reduced solubility in the polar phase, suggesting that these metabolites are not the main active ingredients. Pearson correlation analysis of the MTT values and the relative abundance of each metabolite shows that oleanolic acid and ursolic acid have the highest correlation (Table S11). Moreover, ursolic acid and oleanolic acid are in general abundant in all organs, except for roots, which is in line with the anti-tumor activities of the extracts (Figure 4D, Table S10). The amounts of ursolic acid are highest in the highly active stem and flower extracts of the control conditions and long- and short-day samples. In cold and heat-treated plants, the ursolic acid levels are generally lower. Interestingly, LC-MS/MS identified two compounds related to ursolic acid, of which one (ursolic acid-related 2) is elevated in root extracts of cold-treated (8°C) plants.

To better understand the molecular basis for the different physiological and metabolomic phenotypes of roots, leaves, flowers, and stems we investigated the changes in gene expression. To this end, we produced 42 RNA-seq samples from control leaves, stems, flowers, and roots, and leaves and roots from five stress conditions, cold at 8°C and 15°C, heat, and two high light stresses (Figure 4D). Since the genes of the ursolic and oleanolic acid biosynthesis pathway are characterized in other plants ^48,49^, we carried out homology searches and expression profile analyses of the predicted genes and found two genes that putatively belong to the ursolic acid pathway: *Olcov1051885* and *Olcov1029517. Olcov1051885P1* belongs to the Cytochrome P450 superfamily and is homologous to *Arabidopsis AT5G36130*. *AT5G36130* (of the CYP716A12 family) carries out the last step of ursolate biosynthesis which is the oxidation of α-amyrin to ursolic acid ^50^. The formation of α-amyrin in this pathway is catalyzed by an oxidosqualene cyclase (α-amyrin synthase). We found *Olcov1029517*, a squalene cyclase, that also coexpresses with *Olcov1051885* (PCC=0.94, Table S12). Gene expression of both genes is highest in stems, leaves, and flowers but absent in roots in the control (Figure 4D), which correlates with ursolic acid metabolite levels and MTT activities. In leaves of plants treated with cold, heat, and high light, ursolic acid biosynthesis is decreased along with lower ursolic acid levels (Figure 4E). Interestingly, both genes show expression in roots of cold-treated (8°C) plants, a tissue with elevated anti-tumor activity, and an increase of an ursolic acid-related 2 compound (Figure 4D). Differential gene expression analysis revealed that 3,888 and 5,240 genes are significantly (adjusted p-value < 0.05) up- and down-regulated at 8°C, respectively (Table S13). Moreover, 12 and 28 genes associated with terpene biosynthesis are up- and down-regulated, respectively. Both genes associated with ursolic biosynthesis (*Olcov1051885* and *Olcov1029517*) are significantly upregulated (Table S13) under cold 8°C conditions, but ursolic acid levels were not visibly upregulated in roots (Figure 4D). Conversely, an ursolic acid-related 2 compound is strongly upregulated in roots at 8°C, suggesting that this unknown compound is responsible for the gain of activity in roots at 8°C.

### Ursolic acid causes mitotic catastrophe and cell death in SKBR3

Ursolic acid has been reported to possess a wide range of anti-cancer activities ^51^, such as inhibition of Akt and promotion of autophagy ^52^, a decrease of ATP production ^53^, and activation of adenosine monophosphate-activated protein kinase (AMPK) ^54^. Since many anti-tumor drugs interfere with the cell cycle progression, we set out to study the effect of different concentrations (12.5, 15, 20, 25, and 30µM) of ursolic acid on the cell cycle progression over 24 hours (Figure 5A) and 48 hours (Figure 5B). At lower concentrations at 24 hours of treatment, the cells accumulate at G0/G1 and S phases, while at higher concentrations, the cells no longer accumulate at G0/G1, but instead at G2/M and sub-G1 (sub-G1 represents cells that underwent apoptosis). After 48 hours, we observed more cells in the sub-G1 phase at all concentrations (Figure 5B), indicating that a longer treatment causes more cell death. To track how the ursolic acid influences cell division, we performed time-lapse imaging for 72 hours. In the control samples without ursolic acid treatment, SKBR3 cells are able to undergo at least two rounds of cell division (Figure 5C, Video 1). However, 12.5-20µM of ursolic acid caused delays of the first or second rounds of mitosis (Figure S11), while 20-30µM caused cell death (Figure 5D, Figure S11, video 3).

**Figure 5.**
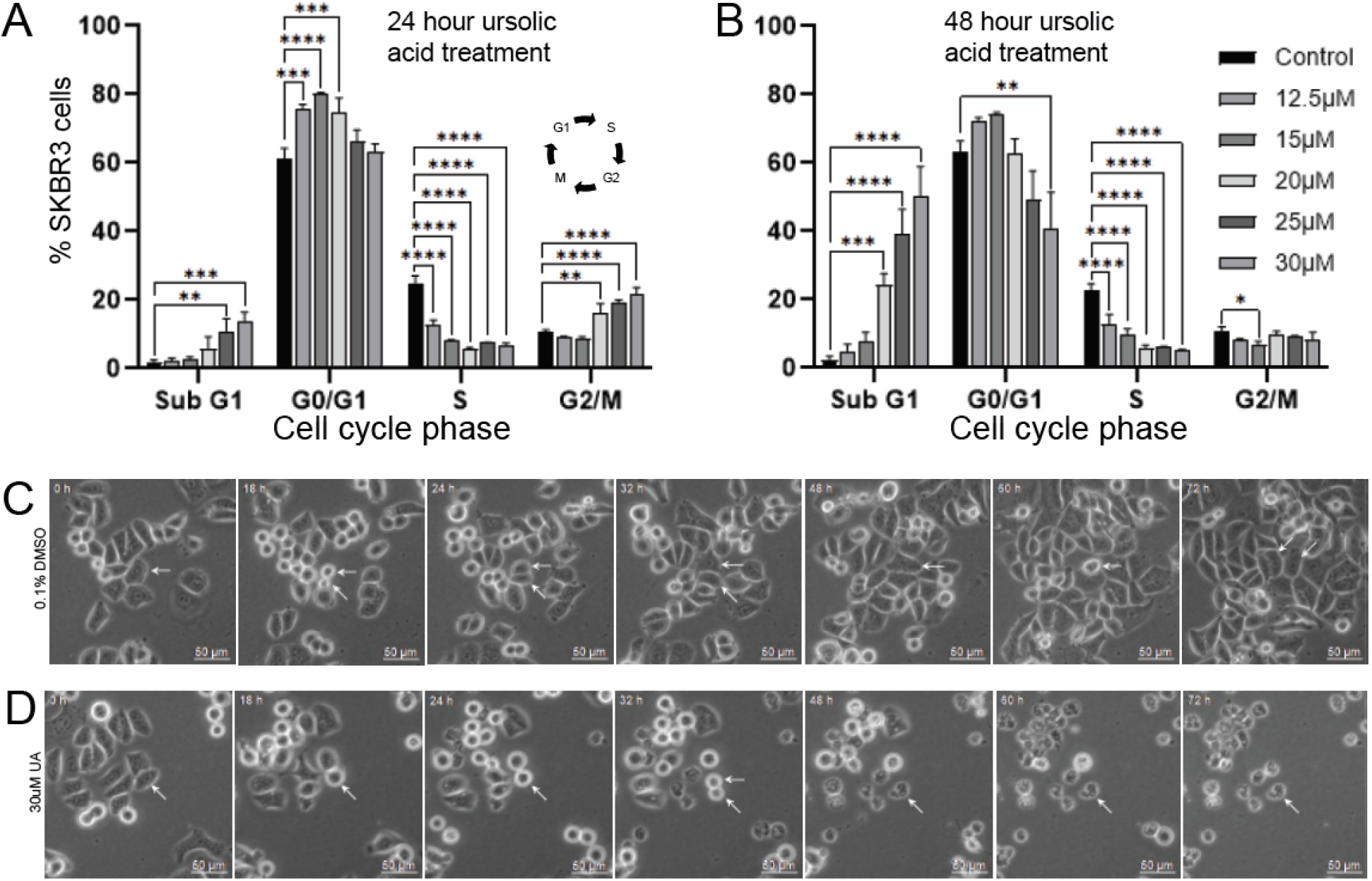
Microscopic analysis of ursolic acid and SKBR3. A) Cell cycle progression analysis of SKBR3 by fluorescence-activated cell sorting (FACS) of DMSO control (black bar) or ursolic acid-treated cells (shades of gray represent ursolic acid concentrations). Each bar represents the mean ± standard deviation from data obtained from three independent sets, where statistical significance was derived through one-way ANOVA followed by Bonferroni’s multiple comparison test (** p < 0.01, *** p < 0.001, and **** p < 0.0001). B) FACS analysis of 48-hour treated cells. C) Phase-contrast images of SKBR3 cells treated with DMSO control over 72 hours. The white arrows are used to indicate a dividing cell (18h). A daughter cell undergoes another division at 60-72 hours. D) Phase contrast image of a cell that undergoes division (32h) and apoptosis (48h).

The results suggest that at high concentrations, ursolic acid may inhibit the mitotic machinery, thereby causing an arrest in mitosis and cell death. At lower concentrations, cells can escape from the mitotic arrest but accumulate cellular damage. Due to the damage in the previous round of mitosis, the cells experience delays in subsequent rounds of cell division, which explains the higher percentage of cells in the G0/G1 stage.

### Cellular thermal shift assay (CETSA) identifies putative human protein targets of ursolic acid

Ursolic acid shows a multi-pronged anti-cancer activity ^51,55^, but its potential protein binding partners in cancer cells are poorly understood. Cellular thermal shift assay (CETSA) is a method to identify protein targets of drugs ^56,57^, and is based on the discovery that ligand binding to a protein increases its thermostability. CETSA combined with multiplexed mass spectrometry (MS-CETSA) represents a powerful untargeted drug-target identification methodology enabling monitoring of the entire detectable proteome for changes in protein thermostability under drug treatment. CETSA-MS does not require prior information on drug type or its mechanism of action ^58^, and its label-free nature enables direct testing of standard unmodified compounds without the need to use expensive drug probes or the generation of reporter cell lines.

To identify protein targets of ursolic acid, we leveraged the multidimensional IsoThermal Drug-dose Response (ITDR) variant of MS-CETSA, where SKBR3 cell lysate was treated with a range of ursolic acid concentrations (10nM to 21.89uM) and exposed to different heat challenge temperatures (50°C, 55°C, 60°C) or a non-denaturing control (37°C) to destabilize susceptible protein subsets. The resulting soluble protein fractions were then analyzed by quantitative mass spectrometry to identify proteins exhibiting drug dose-dependent change in stability, suggestive of direct drug-protein interaction. The MS analysis detected 11,638 protein profiles (Table S14), capturing abundances of 4,356 proteins in at least one thermal challenge and the 37°C reference condition. Of these, 31 proteins passed a higher confidence threshold (3* median absolute deviation in the area under the curve normalized by the reference control, R^2^>0.8).

Among the 31 recorded drug-induced changes in stability, 6 proteins exhibited strong drug-dose dependent stabilizations, expected upon high-affinity drug-ligand interaction, and represent high confidence drug-binding candidates (Figure 6B). The proteins are SPG21 (Maspardin), HIBCH (3−hydroxyisobutyryl−CoA hydrolase, mitochondrial), DECR1 (2,4-dienoyl-CoA reductase [(3E)-enoyl-CoA-producing], mitochondrial), RPLP1 (60S acidic ribosomal protein P1), CWF19-L2 (CWF19−like protein 2) and NIPSNAP (NipSnap homolog 1). The remaining proteins identified by CETSA exhibited only modest positive or negative shifts in their abundance profiles and could represent additional lower confidence off-targets (Figure 6A).

**Figure 6.**
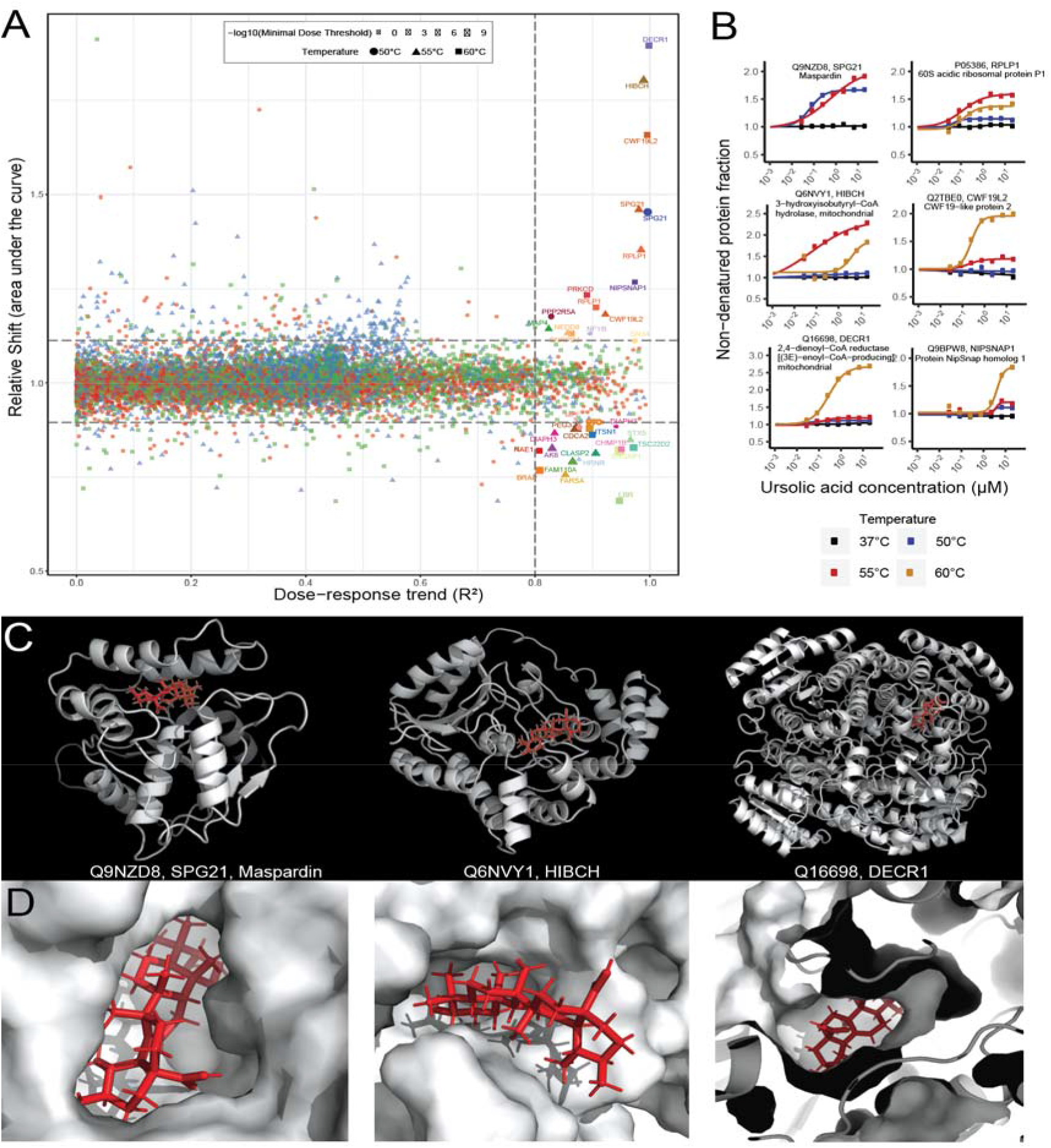
CETSA analysis and reverse docking. A) Distribution of protein stabilization as a function of R^2^ value (that quantifies the adherence of protein stabilization profile to the dose-response trend) against ΔAUC (AUC of heat-challenged sample normalized to non-denaturing 37°C control). 3 * MAD of AUC in each dataset and R^2^ = 0.8 cutoffs are indicated on each graph by dashed lines. Proteins above the dashed thresholds are indicated by names. B) Protein stabilization curves of the top six candidate proteins. The y-axis shows the extent of stabilization, depicted as remaining soluble protein level after thermal challenge relative to no-drug control, while the x-axis shows drug gradients. The non-denaturing 37°C control condition is plotted in black. C) Atomic model of Maspardin, HIBCH, and DECR1. The protein is colored white, while ursolic acid is colored red. D) Zoom in on ursolic acid bound to the respective protein.

To investigate the binding sites of ursolic acid among the identified candidate-binding partners, we performed a reverse docking analysis using ursolic acid and the available protein structures obtained from experimental evidence for DECR1 or AlphaFold predictions for the remaining five proteins. Ursolic acid was able to favorably bind to Maspardin (−13.0 kcal/mol), HIBCH (−8.9 kcal/mol), and DECR1 (−10.6 kcal/mol)(Figure 5D, PyMol session file available as Supplemental Data 1), but not for the other 3 proteins. Interestingly, for DECR1, the docking analysis shows that ursolic acid is potentially able to replace the natural ligands (NADP nicotiamide-adenine-dinucleotide phosphate and hexanoyl-coenzyme A, Figure S12), likely inhibiting the activity of the enzyme.

## Discussion

To identify an *Oldenlandia* plant with potent anti-cancer activities, we screened 11 plants comprising *O. corymbosa*, *O. tenelliflora* and *O. biflora*, and observed that *O. corymbosa* showed specific activity in two breast (SKBR3, MB231) and gastric (SNU484) cancer cell lines, in contrast to the other two plant species (Figure 1). Interestingly, HepG2 and MKN28 were not affected by any of the extracts. As the molecular basis of the cancer cell lines can be different (e.g., mutations in receptors, transcription factors, copy number variations of oncogenes) ^59,60^, some cancer cell lines may need higher concentrations of active compounds to be affected. Interestingly, most other herb garden plants showed either no activity or strong activity on all three cell lines at the concentration used, suggesting that *O. corymbosa* shows a more specific activity to the characteristics of the SKBR3 breast cancer line (Figure S2).

To establish a genomic resource for the *Oldenlandia* genus, and to identify the biosynthetic pathways of the anti-cancer metabolite, we produced a nearly chromosome-scale assembly comprising of 14 contigs. We hypothesize that the high contiguity of the assembly is due to the extremely low heterozygosity of the genome (Table 1), which made the genome assembly easier. The low heterozygosity is comparable to that reported for the homozygous line *Arabidopsis thaliana* Col-0 (heterozygosity of 0.001%)^61^Electron microscopy analysis revealed that the flowers are sealed by a ring of pubescent hairs (Figure 2B), effectively forcing self-pollination (autogamy). Autogamy is thought to be reproductively beneficial in an adverse environment, as there is no need to invest energy in costly pollinating agents and guarantees a greater fruit set ^62,63^. The small, homozygous genome, no evidence of recent whole-genome duplications, short generation time (flowers appear 19 days after germination, Figure 1), and similar experimental procedures to *Arabidopsis* established (Figure S1), make *O. corymbosa* an attractive model for the *Oldenlandia* genus.

It is known that *O. corymbosa* contains iridoids, flavonoids, saponins, triterpenes, phenols, terpenoids, sterols, and anthraquinones ^64,65^. To identify the compounds conferring the anti-cancer activity against SKBR3, we performed activity-guided fractionation, NMR, and high-resolution mass spectrometry, which revealed ursolic acid, oleanolic acid, pheophorbide L, phytol, and lutein in the active fractions. Ursolic and its analog oleanolic acid are found in apple peels and in many herbs ^66^, while pheophorbide L and phytol are produced during chlorophyll a degradation ^67^. Lutein is a xanthophyll carotenoid and antioxidant and has recently been implicated as an anti-tumor agent ^68^. Lutein inhibited the viability of MCF-7 and MDA-MB-231 cells by inducing apoptosis resulting in elevated caspase-3 activity and downregulated expression of Bcl-2 ^68^. Phytol is an emerging candidate with anti-tumor activity, induced genotoxicity, and apoptosis in breast cancer cells. Furthermore, it showed a DNA damage repair capacity in mouse lymphocytes ^69^. Thus, it is likely that the anti-cancer effect of *O. corymbosa* is due to the synergistic action of these compounds.

A main active compound found in both *O. diffusa* and *O. corymbosa* is ursolic acid (Li et al. 2015; Chen et al. 2016). This pentacyclic triterpene shows a range of effects, including anti-tumor, anti-inflammatory, anti-oxidative, antiphytol-diabetic, and pro-apoptotic (Iqbal et al. 2018; Seo et al. 2018; Feng and Su 2019; Sun et al. 2020). The IC50 value of ursolic acid against SKBR3 showed a modest activity of 6.81µM (Table 2), but its activity was more potent than the structurally-similar oleanolic acid (109.7µM) and betulinic acid (66.65µM). This indicates that the relatively minor modification to the compound can have dramatic effects on its efficacy, which makes ursolic acid-like compounds attractive targets for modification, by e.g., synthetic biology approaches ^70^.

Plants produce a wide range of specialized metabolites to better acclimate to changing environments, such as changes in temperature, light, humidity, and developmental processes ^71,72^. The specialized metabolites can serve as toxic, repellant compounds against herbivores, or may be important for functional and structural stabilization through signaling pathways ^47,73^. Thus, amounts of active compounds can be induced by stress. We show that *O. corymbosa* has strong anti-tumor activities in above-ground organs. Interestingly, we observed that the normally inactive roots gain activity in plants grown at 8°C and that this activity is likely caused by an ursolic acid-like compound accumulating in roots (Figure 4D). This is corroborated by the expression of a gene encoding a cytochrome P450 protein (*Olcov1051885*) which is predicted to carry out the last step in the ursolic acid biosynthesis. Co-expressed with this gene is *Olcov1029517*, a putative squalene cyclase. CYP716A (subfamily cytochrome P450) proteins and squalene cyclases are known to be key genes in triterpene biosynthesis in Medicago (Fukushima et al. 2011), apple ^74^, mints ^49^, and Arabidopsis ^75^. Both genes are highly expressed in the stem tissue of the control, correlating with high ursolic acid amounts and strong anti-tumor activity (Figure 4D). Interestingly, our results show that genes related to terpene biosynthesis are upregulated in roots at 8°C. This data suggests that the ursolic acid-like compound found in col-treated roots is synthesized in the roots rather than transported to the roots from above-ground tissues.

Ursolic acid is mentioned in more than 2,600 articles at the time of writing, and it has been observed to have many health-promoting activities. The compound has a wide range of anti-tumor activities and directly affects the hallmark of cancer growth, which is the metabolic reprogramming of cancer cells to sustain anabolic growth ^76^. Ursolic acid can modulate the expression and function of mitochondria-associated enzymes, which promote apoptosis and lead to anti-proliferative effects in various models in vivo and in vitro (ref). In three breast cancer cell lines with different growth factor receptor status (MCF7, MB-231, SKBR3), which we also used in our study, ursolic acid downregulates the Akt signaling, and this inhibition affects glycolysis and markedly decreased levels of HK2, pyruvate kinase M2, ATP and lactate (Lewinska et al. 2017). Additionally, ursolic acid and betulinic acid (5 and 10 µM) caused p21-mediated G0/G1 cell cycle arrest and senescence accompanied by oxidative stress and DNA damage. 20 µM ursolic acid-induced apoptosis was achieved by targeting the glycolytic pathway and autophagy in breast cancer cells ^52^. Similarly, another study shows that ursolic acid derivatives induce cell cycle arrest at the G2/mitotic (M) phase by competing with glucose for the binding site at glucokinase (Wang et al. 2014). These results are in line with our observations for SKBR3 cells, where we observed a concentration- and time-dependent accumulation of cells at G0/G1 and S phases (low concentrations) or G2/M and sub-G1 (high concentrations), with more cells accumulating at sub-G1 phase (representing cell death) at 48 hours (Figure 5). Thus, we hypothesize that ursolic acid causes mitotic cycle arrest.

Despite the numerous studies, the direct protein binding partner of ursolic acid is unknown. To address this, we performed CETSA-MS, and identified six high-confidence proteins. Three of them (HIBCH, DECR1, RPLP1) have been implicated in cancer. HIBCH is involved in L-valine degradation, and blocking HIBCH localization to mitochondria resulted in decreased colon cancer growth and increased autophagy *in vivo* and *in vitro* ^77^. HIBCH has also been reported to be increased in patients with prostate cancer and MDA-MB-231 breast cancer ^78,79^. DECR1 is a short-chain dehydrogenase involved in redox homeostasis by controlling the balance between saturated and unsaturated phospholipids. *In vivo*, DECR1 deletion impairs lipid metabolism and reduces tumor growth in prostate cancer ^80^. In cervical cancer high levels of DECR1 enhance lipolysis and the release of fatty acid energy stores to support cancer cell growth ^81^. In contrast, *DECR1* was proposed to act as a tumor suppressor in HER2-positive breast cancer, however, DECR1 overexpression was shown to protect cancer cells from glucose withdrawal-induced apoptosis ^82^. The 60S acidic ribosomal protein P1 (*RPLP1*) plays an important role in the elongation step of protein synthesis and promotes tumor metastasis in breast cancer patients, as well as in hepatocellular carcinoma cells and several gynecologic tumors ^83–86^.

The remaining three proteins have diverse roles. The first protein, Maspardin (SPG21), may play a role as a negative regulatory factor in CD4-dependent T-cell activation as it co-localizes with CD4 on endosomal/trans-Golgi network ^87^, which is in line with ursolic acid affecting T-cell activation ^88^. The second protein, CWF19-like protein 2, is a cell cycle control factor located in the nucleus predicted to be involved in mRNA splicing via spliceosome. It appeared in a screen for chromosomal deletion sites in breast carcinomas together with other cell-cycle regulation proteins (JMY, PTPRN2)^89^. In addition, it might interact with BRCA1 ^90^ and BRCA2 ^91^. The third protein, NIPSNAP homolog 1, is involved in removal of damaged mitochondria and was shown to be upregulated upon treatment with anti-tumor drugs ^92–94^.

Taken together, HIBCH, DECR1, and RPLP1 constitute prime targets for their role in the ursolic acid-mediated activity.

## Conclusion

We present the first high-quality genome for the *Oldenlandia* genus and characterize the mode of action of the main active compound - ursolic acid. We envision that the genome will be valuable for researchers working on the medicinal properties of the *Oldenlandia* species, while the mode of action of ursolic acid will spur further development of this valuable compound.

## Material and Methods

### Establishing *Oldenlandia* plants in the laboratory

*Oldenlandia* plants (*O. corymbosa, O. tenelliflora* and *O. biflora*) were collected in the field in the Bedok area of Singapore. Ten individual plants (M1, M2, M5, M6, M7, M9, M11-14) were transferred to the lab and naturally propagated by autogamy. Line 15 (*O. corymbosa*) was collected from a private residential area. Plantaflor potting soil, vermiculite, and perlite were used as a mixture (7:2:1). Standard conditions in the growth chamber were 28°C, 50-60% humidity and 12h light (∼200 μmol·m^-2^·s^-1^).

Plants used for the initial MTS assays (Figure 1D) were grown at 23°C, 50-60% humidity and 12h light conditions (150 μmol·m^-2^·s^-1^). MTS assays of *O. corymbosa* compared to medicinal herb garden plants (Figure 1S) were done on five F1 plants of M15 grown outside at 28-35°C, 60-80% humidity and 12h variable light conditions (150-1500 μmol·m^-2^·s^-1^). Abiotic stress experiments were done using F2 plants of M15. We grew plants in soil under normal conditions (200 μmol·m^-2^·s^-1^, 28°C, 12h light) until DAG 23 (±2 days, until 4^th^ node was developed, Figure S9A) and then transferred them to eight different stress conditions for 7 or 9 days. The 7-day long treatments included short day (8h light/16h dark), long day (20h light/4h dark), cold (8°C and 15°C), and heat (40°C). The 9-day long treatments were drought (no watering) and two high light conditions (1000 and 2000 μmol·m^-2^·s^-1^). Drought plants were left unwatered for 9 days until leaves were rolled in. One plant was watered on the day of harvest and fully recovered the next day, thus showing that plants were still viable at the time of harvest (not shown). Flowers, leaves, and stems were harvested and directly frozen in liquid nitrogen. Roots were rinsed in water for about 5-8 min to remove soil and vermiculite, dried with paper and then frozen in liquid nitrogen.

### RNA isolation and sequencing

An RNA tissue atlas for genome annotation was prepared from flowers, leaves, stems and roots of M15 plants grown outside under low light conditions using Spectrum^TM^ Plant Total RNA Kit (Sigma). For differential gene expression analyses of secondary metabolite genes, RNA was extracted from stressed and control plants (three biological replicates each) shown in Figure 4A using Spectrum^TM^ Plant Total RNA Kit (Sigma). Quality control of all the extracted RNA (triplicates for each condition) was done by Novogene (Singapore) using Nanodrop and agarose gel electrophoresis (for purity and integrity) before sample quantitation and further analyses of integrity (Agilent 2100 Bioanalyzer). Library type was a eukaryotic directional mRNA library. Library construction from total RNA including eukaryotic mRNA enrichment by oligo(dT) beads, library size selection, and PCR enrichment was performed by Novogene using NEBNext® Ultra™ II Directional RNA Library Prep Kit for Illumina®. The libraries for the RNA tissue atlas were then sequenced with Illumina Hiseq-4000, paired-end sequencing at 150 base pairs and at sequencing depths of ∼51 million reads per sample (15Gb/sample). The libraries for the abiotic stress experiments were sequenced with NovaSeq-6000, paired-end sequencing at 150 base pairs and at sequencing depths of ∼20 million reads per sample (6Gb/sample).

### DNA Extraction and Illumina Sequencing

Genomic DNA was isolated from 100 mg (F.W.) of leaf tissues from M15 plants grown outside under low light conditions using the illustra Nucleon Phytopure Genomic DNA Extraction Kit (GE Healthcare) after grinding the leaves in liquid nitrogen to a fine powder. A total amount 5.85 ug (A260/280: 1.84; A260/230: 1.36) was sent for Illumina sequencing at Novogene AIT (Singapore). Library type was a 350 bp insert DNA library (WGS). The libraries were then sequenced with Illumina HiSeq-4000, paired-end sequencing at 150 base pairs and at sequencing depths of ∼102 million reads per sample (30Gb/sample).

### High Molecular Weight DNA Extraction and Oxford Nanopore sequencing

Leaves and stems of the clonally propagated F1 plants of M15 (*Oldenlandia corymbosa*) were harvested, cleaned and cut into smaller pieces, and care was taken to exclude flowers and fruits and seeds. Plant tissue was flash-frozen in liquid nitrogen and stored at −80°C prior to High Molecular Weight (HMW) DNA extraction. For each extraction, 10 g of plant tissue was physically homogenized in liquid nitrogen using a mortar and pestle. The BioNano Nuclear Isolation protocol was followed, in which the homogenized plant tissue was first dissolved in Isolation Buffer with Triton X-100 and β-mercaptoethanol (IBTB), followed by filtering to remove undissolved plant material, and then centrifugation was carried out to pellet the nuclei. Isolation Buffer (IB) consisting of 15 mM Tris, 10mM EDTA, 130mM KCl, 20 mM NaCl, 8% PVP-10, pH 9.4 was prepared, with 0.025% (m/v) spermine and 0.035% (m/v) spermidine added and filter-sterilized before use. 0.1% Triton X-100 and 7.5% β-mercaptoethanol were next added to the IB, and the resulting IBTB buffer was added to the homogenized plant tissue and mixed on ice. Cell strainers (100 μm and 40 μm) were used to remove the undissolved plant tissue. Triton X-100 was added to the cell suspension to facilitate the disruption of the cell membrane and the samples were mixed thoroughly before being spun down at 2000 *g* for 10 min to pellet the nuclei. The pelleted nuclei were resuspended in cetyl trimethylammonium bromide (CTAB) solution for DNA extraction following the protocol described in Michael et al. (2018) ^95^. The resulting DNA was further purified using the Qiagen Genomic Tip 500/G. The DNA pellet from the CTAB DNA extraction was solubilized in Buffer G2 from the Qiagen Genomic Tip kit, along with RNase A (2 μl/ml of buffer) and proteinase K (10 μl/ml of buffer) for further removal of contaminating RNA and proteins. The rest of the Qiagen Genomic Tip protocol was followed as per the manufacturer’s instructions. The final DNA extract was quantified using Nanodrop and the DNA integrity was checked using the Genomic ScreenTape (Agilent). Finally, HMW DNA was size-selected and sequenced as previously described ^96^. In brief, DNA was size-selected using the Circulomics Short-Read Eliminator XL Kit and a sequencing library was prepared using the Oxford Nanopore LSK-110 ligation sequencing Kit. The library was sequenced using a PromethION using 9.4.1 cells. Basecalling of the ONT read data was done using guppy basecaller v3.2.2, which yielded ∼2.7 million reads.

### Genome assembly

To assemble the genome of *O. corymbosa* we used long reads from Oxford Nanopore sequencing and short reads from Illumina sequencing. ONT reads were processed by removing adapters using porechop v0.2.3 (https://github.com/rrwick/Porechop) with default settings, removing low quality (<85 q) and short reads (<3,000bp) using filtlong v0.2.0 (https://github.com/rrwick/Filtlong), and error-corrected using Canu v1.9 ^97^. For Illumina reads, low quality sequences and adapters were trimmed using Trimmomatic v0.39 ^98^ with default parameters. To estimate the genome size of *O. corymbosa*, jellyfish v2.3.0 ^99^ was run on the Illumina reads with the canonical k-mer (−C) option and k-mer size of 21. Then GenomeScope v2.0 ^100^ was used to analyze the k-mer distribution, which resulted in a haploid genome size of 283.8Mbp. ONT corrected trimmed reads were assembled using Canu v1.9 ^97^ with the estimated genome size. ONT corrected trimmed reads were assembled using Canu v1.9 with the estimated genome size. Then, contigs were polished for two rounds using RACON v1.4.3 ^101^ with the alignments generated by MiniMap2 v2.17 ^102^ using the ONT reads, and Pilon v1.24 ^103^ with the alignments generated by BWA v0.7.17 ^104^ using the Illumina reads. Duplicate haplotypes were removed using Purge Haplotigs ^105^ with default parameters and read dept thresholds of 5, 45, and 190. Finally, two rounds of polishing were done using RACON and Pilon as described before.

### Genome annotation

The *de novo* repeat library of *O. corymbosa* was constructed following the Repeat Library Construction Advanced pipeline ^106^, which uses MITE-Hunter ^107^, LTRdigest ^108^, LTR_harvest ^109^ (available in genome tools v1.6.1 ^110^) and Repeatmodeler v2.0.1 ^111^. This new library was combined with the Dfam v3.5 database ^112^ to identify Transposable Element. The genome of *O. corymbosa* was masked using both libraries with RepeatMasker v4.1.2-p1 ^113^. Gene prediction was performed by combining transcript alignments, protein alignments and *ab initio* gene predictions ^114^. First, the RNA-seq reads were trimmed using Trimmomatic v0.39 ^98^ with default parameters. Then, a *de novo* transcriptome assembly was obtained with Trinity v2.11.0 ^115^ and non-redundant transcripts were obtained with PASA v2.5.1 ^116^. After, we detected coding regions in the transcript using TransDecoder from the PASA package. Second, we downloaded the proteomes of two Rubiaceae (*Coffea canephora* and *Leptodermis oblonga*) from NCBI database and aligned to the genome using SPALN v2.4.6 ^117^. Third, the RNA-seq reads were aligned to the genome with STAR v2.7.8a ^118^ and hints were generated with bam2hints from the Augustus package v3.2.3 ^119^. Then, ab initio gene predictions were performed with Augustus v3.2.3, GeneID v1.4 ^120^, and GeneMark-ES v4 ^121^ with and without hints. Finally, all the predictions were combined using EVidenceModeler-1.1.1 ^116^. Functional annotation was performed using InterProScan v5.55-88.0 ^122^ and Mercator4 v4.0 ^123^. Visualization of the genome characteristics was performed with Circos v0.69-8 ^124^.

### Plastid genome assembly and annotation

The plastid genome was recovered from the Illumina reads using GetOrganelle v1.7.5 ^125^ and annotated using GeSeq ^126^. Then, the genome map was drawn with OGDRAW ^127^

### Genome analysis

Assembly completeness was estimated in three ways. First, gene completeness was determined by running BUSCO v5.3.1 ^128^ using the eudicots_odb10 database. Second, a comparison of k-mers present in the assembly and in the Illumina reads was performed using Merqury ^129^ with k-mer length of 21. Third, telomere repeats (‘TTTAGGG’) were searched in the contigs. Then adjacent regions were joined, and regions larger than 100 bp were included as predicted telomere regions.

To detect heterozygous positions in the genome, Illumina reads were mapped against the genome using BWA v0.7.17-r1188 ^104^, and SNPs were identified with GATK HaplotypeCaller v4.2.5.0 ^130^, setting ploidy to 2, and using thresholds for mapping quality (MQ > 40), quality by depth (QD > 2), Fisher strand bias (FS < 60), mapping quality rank sum test (MQRankSum > − 12.5), read pos rank sum test (ReadPosRankSum > − 8), strand odds ratio (SOR < 3), read depth of coverage (DP ≥ 20), and allelic depth (AD ≥ 5).

### Prediction of orthogroups and species tree reconstruction

To study the evolution of *O. Corymbosa* in the context of other nine plant species, orthogroups (orthologous gene groups) were predicted using Orthofinder v2.5.4 ^131^ with Diamond v2.0.5.143 ^132^ as sequence aligner. A total of 807 single-copy orthogroups were selected and aligned using MAFFT v7.453 ^133^ individually and then concatenated. The species tree was reconstructed using the amino acid substitution model LG implemented in RAxML v8.2.12 ^134^ and 100 bootstrap replicates. Additionally, the assembled plastid genome of *O. corymbosa* was aligned with other five genomes (*O. corymbosa*, *O. diffusa*, *O. brachypoda*, *Hedyotis ovata*, and *C. arabica*) using MAFFT v7.453 ^133^ and a species tree was reconstructed as described above using RAxML v8.2.12 and the GTR model.

### RNA seq analysis

To obtain transcripts abundances of the RNA-seq samples under different conditions, the reads were mapped to the coding sequence (CDS) of *O. corymbosa* with Kallisto v.0.46.1 ^135^. All samples had >8M reads mapped. For root and stem samples, differential expression analysis was performed with the package DESeq2 ^136^.

### MTT, Metabolic, and RNA profiling

To detect which of the candidate metabolites is responsible of the anti-cancer activity, Pearson correlation coefficient (PCC) of the percentage of cell viability (MTT assay) profiles was calculated against the relative abundance of Ursolic acid, Oleanolic Acid, Pheophorbide a, and two compounds related to Ursolic acid.

To study the genes associated with the metabolic pathway of ursolic acid, candidate genes were searched in other species in the PlantCyc database ^48^. Then homology searches in the O. Corymbosa were done using Blast and the orthogroup information. Additionally, the profiles of expression of the candidate genes were analysed and compared with the metabolic profiles. Finally, gene coexpression analysis was performed by calculating the PCC against all the proteomes and the 50 best hits were kept.

### Anti-cancer activity, Metabolic, and RNA profiling

To detect which of the candidate metabolites is responsible of the anti-cancer activity, Pearson correlation coefficient (PCC) of the percentage of cell viability (MTT assay) profiles was calculated against the relative abundance of Ursolic acid, Oleanolic Acid, Pheophorbide a, and two compounds related to Ursolic acid.

To study the genes associated with the metabolic pathway of ursolic acid, candidate genes were searched in other species in the PlantCyc database ^48^. Then homology searches in the *O. corymbosa* were done using Blast and the orthogroup information. Additionally, the profiles of expression of the candidate genes were analysed and compared with the metabolic profiles. Finally, gene coexpression analysis was performed by calculating the PCC against all the proteomes and the 50 best hits were kept.

### Metabolite extraction and LC-MS/MS analysis

For the abiotic stress experiment (Fig 3B) four major tissues were sampled. Flower tissue (open and closed flowers and green seed pods) was collected and frozen in liquid nitrogen. Leaves of >2cm length were cut, rinsed in dH2O, pat dry on a paper tissue and flash frozen. Smaller leaves and apical tissue were spared. Stems were cut and flash frozen. Roots were carefully rinsed for 5-8 min in water to remove soil particles, dried with paper tissue and flash frozen. All samples were ground using a mortar and pestle, the frozen powder was aliquoted and stored in −80°C until use.

For metabolite extraction, 5μl/mg F.W. of extraction buffer (80% MeOH in water, HPLC-MS grade) was added to 100 mg of frozen powder together with 3 stainless steel beads. The samples were homogenized in a Qiagen PowerLyser for 2 min at 2000 rpm. Beads were removed and samples were shaken at 1500 rpm for 10 min at 20°C in the dark followed by centrifugation at 12,000 rpm for 10 min. After transfer of supernatant, the pellets were re-extracted twice and the collected supernatants were spun again, aliquoted and dried in a cooling speed vac at 30°C. Pellets were stored in −80°C. For MTT analyses, pellets were dissolved at a concentration of 2 μl DMSO/mg frozen powder. Samples were centrifuged at 13,000 rpm for 10 min and the SUP was filtered through 0.2 μm nylon filters (Sigma CLS8169) before use.

For Figure 1D 100 mg of fresh leaf sample was homogenized in 100% MeOH in a PowerLyser (at 2500 rpm for 30 sec) and centrifuged at 12,000 rpm for 10 min. The pellet was re-extracted 2 times. For Figure S1 100 mg of lyophilised plant material was ground to fine powder in a PowerLyser and then extracted in 1 ml 100% MeOH for 2 hours on a shaker at room temperature. After centrifugation at 12,000 rpm for 10 min, the SUP was left to dry in a fume hood. After estimating the weights of the pellets they were dissolved in 50-200 μl DMSO and used at extract concentrations of 0.5-2.0 ul/mg

For LC-MS/MS analysis, the polar fraction was dried under vacuum, and the residue was resuspended in 250 μL of LC-grade water. After sonication of samples for 10 min in an ice-cooled sonicator bath, tubes were centrifuged for 15 min at full speed (>12,000 rpm). The supernatant (100 μL)—without any sediment from the bottom of the tube—was transferred to LC tubes for analysis. The samples were run on a UPLC-LC-MS machine as described previously ^137^. In brief, the UPLC system was equipped with an HSS T3 C18 reverse-phase column (100 × 2.1 mm internal diameter, 1.8 μm particle size; Waters) that was operated at a temperature of 40°C. The mobile phases consisted of 0.1% formic acid in water (solvent A) and 0.1% formic acid in acetonitrile (solvent B). The flow rate of the mobile phase was 400 μL/min, and 2 μL of sample was loaded per injection. The UPLC instrument was connected to an Exactive Orbitrap-focus (Thermo Fisher Scientific) via a heated electrospray source (Thermo Fisher Scientific). The spectra were recorded using full-scan positive and negative ion-detection mode, covering a mass range from m/z 100 to 1,500. The resolution was set to 70,000, and the maximum scan time was set to 250 ms. The sheath gas was set to a value of 60 while the auxiliary gas was set to 35. The transfer capillary temperature was set to 150°C while the heater temperature was adjusted to 300°C. The spray voltage was fixed at 3 kV, with a capillary voltage and a skimmer voltage of 25 V and 15 V, respectively. MS spectra were recorded from minutes 0 to 19 of the UPLC gradient. Processing of chromatograms, peak detection, and integration were performed using RefinerMS (version 5.3; GeneData). Metabolite identification and annotation were performed using standard compounds, tandem MS (MS/MS) fragmentation, metabolomics databases and an in-house reference compound library.

### Cell cultures

MCF7 and MDA-MB231 cells were maintained in RPMI-1640 (Thermo Scientific Hyclone, South Logan, UT, USA) containing 10% FBS (Hyclone) and 100U penicillin/streptomycin (Biological Industries, Israel). SKBR3 was maintained in DMEM (Hyclone) containing 10% FBS and 100U penicillin/streptomycin.

### Assessing metabolic activity of cancer cells with MTS and MTT assays

For the MTT (3-(4,5-dimethylthiazol-2-yl)-5-(3-carboxymethoxyphenyl)-2-(4- sulfophenyl)-2H-tetrazolium) cell viability assays in Figure 1D and S1 a total of 2,500 cells were seeded per well in triplicate in 96-well plates one day prior to the cell proliferation assay to allow for cell attachment. On the following day, cells were treated with the different extracts or vehicle control at 0.2 µl/100 µl medium for 48 hours at 37°C. The medium was aspirated and fresh medium containing 3-(4,5-dimethylthiazol-2-yl)-5-(3-carboxymethoxyphenyl)-2-(4-sulfophenyl)-2H-tetrazolium (MTS, premixed in 5:1 ratio, Promega) was added to the treated cells. After 1 hr incubation at 37°C in the dark, the optical density was read at 490 nm using a plate reader.

For MTT assays in Figures 3B and Figures 4B-C and 4E around 11,000 SKBR3 cells were seeded per well in DMEM and incubated at 37°C overnight to allow for cell attachment. The following day cells were treated with the different extracts. After 48h incubation at 37°C the media was aspirated and replaced with fresh DMEM media. After addition of 10μl of 3-(4,5-dimethylthiazol-2-yl)-2,5-diphenyl tetrazolium bromide (MTT) solution (5mg/ml, Sigma-Aldrich, USA) the cells were incubated in darkness for another 4 hrs. Formazan crystals were dissolved with a solubilising agent. For Figure 3B and 4B-C 100 μl acidified SDS (10% SDS in 0.01M HCl) was added followed by incubation at 37°C for 4 hours in the dark. Alternatively, the crystals were solubilised by adding 100 μl DMSO and shaking the plate at room temperature for 15-30 minutes (for Figure 4E). Optical density was read at 570 nm using a plate reader.

For Figure 3B one μl of 100 μl DMSO-dissolved methanol extract (F.W. 50 mg powder) was used in the assay. For Figure 4B the pellet of total minus PK3507 was dissolved in 30 μl DMSO and the pellet containing the purified peak PK3507 was dissolved in 10 μl. Dilution curves were prepared for both samples and 4 μl were used in the MTT assay. To assay the 24 fractions (Figure 4C), each fraction was resuspended in 10 μL DMSO and 1 μl was added to the assay. To determine IC_50_ values all fractions were diluted as indicated in Figure 4E and 0.1% DMSO was used for control wells. IC_50_ values of isolated compounds and standards were calculated by GraphPad Prism 4 software. All determinations were done in triplicates.

### Time-lapse imaging of SKBR3 cells treated with ursolic acid

Cells were seeded on a 6-well tissue culture plate, treated with drugs/fractions, and placed on a heat-controlled stage of a Zeiss Axiovert 200M inverted microscope equipped with a 0.3 NA EC Plan-NeoFluar 10x lens (Carl Zeiss, Germany). The temperature was maintained at 37°C and CO2 levels were maintained at 5%. Phase-contrast images were acquired at 15 minutes intervals for 72 hours.

### Cell cycle analysis by flow cytometry

2.75 × 105 SKBR3 cells were seeded on a 6-well tissue culture plate and cultured for 72 hours before treatment. Cells were treated with DMSO (0.1%) and varying concentrations of ursolic acid (12.5-30μM) for 24 and 48 hours. Cells were harvested, washed with PBS and fixed in 70% ice-cold ethanol overnight. Fixed cells were stained in a solution containing 0.1% Triton-X 100, 10μg/ml RNase (Sigma-Aldrich) and 20μg/ml Propidium Iodide (PI)(Sigma-Aldrich). PI fluorescence was detected using BD LSRFortessa X-20 Cell Analyser (Becton, Dickinson & Company). Cell cycle analysis was performed using FlowJo software (Becton, Dickinson & Company). All experiments were performed in triplicates.

### Identification of putative cysteine-rich peptides (cyclotides) with high-resolution LC-MS analysis

Methanolic extracts of *O. corymbosa* with anticancer activity to SK-BR3 (CIII-L2 and L3) were injected into an Orbitrap Elite mass spectrometer (Thermo Scientific Inc., Bremen, Germany) coupled with a Dionex UltiMate 3000 UHPLC (ultra-high-performance liquid chromatography) system (Thermo Scientific Inc., Bremen, Germany) to identify putative cysteine-rich peptides within the molecular weight range of 2000 to 6000 Da. Samples were sprayed using a Michrom’s Thermo CaptiveSpray nanoelectrospray ion source (Bruker-Michrom Inc, Auburn, USA) and separation was performed using a reversed-phase Acclaim PepMap RSL column (75LLμm IDLL×LL15Lcm, 2 μm; Thermo Scientific). The mobile phase was 0.1% formic acid (FA) as eluent A and 90% ACN 0.1% FA as eluent B, with a flow rate of 0.3 μL/min. A 60 min gradient was used for the elution as follows: 3% B for 1 min, 3–35% B over 47 min, 35–50% B over 4 min, 50–80% B over 6 s, 80% for 78 s; then, it was reverted to the initial state over 6 s and maintained for 6.5 min.

### Cyclotide purification by reversed-phased high-performance liquid chromatography (RP-HPLC) fractionation

Methanolic extracts of *O. corymbosa* with anticancer activity to SK-BR3 (CIII-L2 and L3) were fractionated using RP-HPLC (Shimadzu, Japan). A linear gradient of mobile phase A (0.05% TFA in H2O) and mobile phase B [0.05% TFA in acetonitrile (ACN)] was used with the C18 column (250 x 4.6 mm, 5 µm, 300Å; Phenomenex, USA). MALDI-TOF MS (AB Sciex, USA) was used to identify the presence of the 3507 Da compound in the eluted fractions which were then pooled to form the 3507 Da-enriched fractions. The remaining fractions were pooled together to form the 3507-Da-depleted fraction. The combined samples were then lyophilized for storage. For MTT assays, the pellet of total minus the PK3507 was dissolved in 30 μl DMSO and the pellet containing the purified peak PK3507 was dissolved in 10 μl. Dilution curves were prepared for both samples and 4 μl were used in the MTT assay.

### Identification of active metabolites with activity-guided fractionation

Frozen pulverized plant material (100 mg leaves of M15) was extracted with methanol (1 mL) in an ultrasonic bath for 20 mins. The sample was centrifuged and the supernatant was dried under vacuum. The dried extract was reconstituted in methanol and fractionated on an analytical scale HPLC. 23 fractions and crude were submitted for biological testing. Two active fractions (6N and 6P) were observed.

Scale-up fractionation was carried out to obtain more 6N and 6P, and to further purify the major compounds. Oven-dried milled plant leaves and stems (7 g) were extracted twice with dichloromethane / methanol (1:1) in an ultrasonic bath for 20 mins. The filtered extracts were combined and dried under vacuum. The dried extract (405 mg) was reconstituted in methanol and 250 mg was fractionated using a Preparative-HPLC. The fractions isolated were analysed by ^1^H NMR and HR-MS. Four known compounds were isolated, namely ursolic acid (1), pheophorbide L (2), phytol (3) and lutein (4). These compounds were identified by comparison of their spectroscopic data with those reported in the literature ^138–141^.

### Analytical scale fractionation by UPLC/QTOF

Analytical scale fractionation was carried out on an Agilent UPLC1290 coupled with a Quadrupole Time-of-Flight (Q-TOF) system. Separation was carried out with a reversed-phase C18 column (4.6 x 75 mm) at 2 mL/min, using a 20 mins linear gradient with 0.1% formic acid in both solvent A (water) and solvent B (acetonitrile). The typical QTOF operating parameters were as follows: positive ionization mode; sheath gas nitrogen flow, 12 L/min at 295°C; drying gas nitrogen flow, 8 L/min at 275°C; nebulizer pressure, 30 psi; nozzle voltage, 1.5 kV; capillary voltage, 4 kV. Lock masses in positive ion mode: purine ion at m/z 121.0509 and HP-921 ion at m/z 922.0098.

### Scale-up fractionation by Preparative HPLC

Scale-up fractionation was performed on an Agilent HPLC1260 coupled with a 6130 Single Quadrupole (SQ) system. The extract was fractionated by a reversed-phase column C18 (30 x 100 mm) at 48 mL/min, under a gradient condition from 40-100% solvent B in 60 mins. Ursolic acid, pheophorbide L, lutein and phytol were eluted at retention time 36, 42, 53 and 56 mins respectively. Bruker DRX-400 NMR spectrometer with Cryoprobe, and 5-mm BBI probe head equipped with z-gradients was utilized to obtain ^1^H NMR spectra of the compounds.

### Variable Pressure-SEM

The flower was visualized untreated in a variable-pressure scanning electron microscope VP-SEM (Hitachi FlexSEM 1000 II) at 30 Pa and accelerating voltage of 10 kV, using a BSE detector.

### Cellular Thermal Shift Assay

ITDR MS-CETSA was essentially done as in ^58^ with the following specifications. Cancer cell lysate of SKBR-3 was exposed to varying range of drug concentrations (65.67uM - 10nM) and exposed to 4 different heat challenge temperatures 50°C, 55°C, 60°C and 37°C as non-denaturing control. Remaining soluble protein fractions were analysed by quantitative mass spectrometry to identify proteins exhibiting drug-dose dependent change in stability, suggestive of direct drug-protein interaction. During the data analysis step, two points were removed due to substantial deviations in relative protein abundances (the top concentration data point, due to a large number of single-point (likely non-specific) apparent stabilizations/interactions, as well as the no-drug control due to a likely pipetting error). Final protein stability profiles were plotted along the 10 nM-21.89 uM drug concentration gradient.

### Molecular dynamics simulation

The structure models of the proteins, Q9NZD8 Maspardin and Q6NVY1 3−hydroxyisobutyryl−CoA hydrolase, mitochondrial, were build using Alpha Fold2 server ^142^. The structure of Q16698 2,4-dienoyl-CoA reductase [(3E)-enoyl-CoA-producing], mitochondrial, was taken from the PDB 1W6U ^143^. Both proteins and the ligand, ursolic acid, were constructed using the CHARMM-GUI ^144^ tool. Parameters of the proteins were based on the CHARMM36 force field ^145^ and parameters of the ligand were based on the CHARMM General Force Field ^146,147^. The binding pockets of the proteins were identified by PointSite ^148^. The quick vina2 ^149^ was used to dock the ligand to the binding pockets. The best poses were chosen to be the initial structures for the following molecular dynamics simulations by manually checking all the vina docking output. All the topological files were converted to GROMACS ^150^ format using CHARMM-GUI force field converter. The system were solvated with TIP3P ^151^ water molecules and counterions were added to neutralize the system. The three protein-ligand systems were subjected to molecular dynamics (MD) simulation using GROMACS v5.1.2 software ^150^. The LINCS ^152^ algorithm was used to constrain bonds between heavy atoms and hydrogen to enable a time step of 2fs. A 1.2nm cutoff was used for van der Waals interaction and short-range electrostatic interactions calculations, and the Particle Mesh Ewald method was implemented for long-range electrostatic calculations. Simulation temperature was maintained at 300K using a V-rescale thermostat ^153^ and 1bar pressure using Parrinello-Rahman ^154^ barostat. Each simulation lasted 100 ns. The representative structures were taken as the center of the biggest cluster got from the gromos clustering method ^155^.

### Data availability

All the data generated or analyzed in this study are included in this article. The genome assembly, its annotation, and the sequencing data (DNA and RNA) have been deposited in the ENA (European Nucleotide Archive) under the accession number PRJEB50026.

## Acknowledgments

We would like to thank Victor Albert and Charlotte Lindqvist for useful discussions, and the members of the Mutwil lab for feedback.

## Supplemental tables

**Table S1. Collected Oldenlandia species and locations.**

**Table S2. MTT assays testing the anti-cancer activity of several Oldenlandia species.** The top table shows the average anti-cancer activity, while the lower table shows the standard deviation of the average.

**Table S3. Characteristics of the plastid genomes of different Oldenlandia and Hedyotis species.**

**Table S4. Characterization and quantification of repetitive sequences in the O. corymbosa genome.**

**Table S5. List of species used in this study.** The columns in order show the species name, taxon ID, number of proteins, genome size, and the source of the proteome.

**Table S6. MTT assay activities against SKBR3 cancer cell lines for the cyclotide enriched dilutions (3518Da - 3518Da), and cyclotide-depleted diultions (Depleted - Depleted 18:8).** The measurements are taken for three technical replicates.

**Table S7. MTT assay activities against SKBR3 cancer cell lines for the initial fractionation.** The measurements are taken for three technical replicates.

**Table S8. Dilution series for the fractions and standards.** MTT assay activities against SKBR3 cancer cell lines for the four fractions and five standards. The measurements are taken for three technical replicates.

**Table S9. MTT assay activities against SKBR3 cancer cell lines from plants exposed to abiotic stress. T**he columns are: sample ID, the sampled organ, the applied stress, the viability value inferred from the MTT assay.

**Table S10. Relative abundance of ursolic acid, oleanolic acid, and pheophorbide** □ **in the same samples used for the MTT assay.** The values are inferred from LC-MS/MS analysis.

**Table S11. Pearson correlation coefficient (PCC) of the MTT assay profiles versus the relative abundance of five metabolites.**

**Table S12. Pearson correlation coefficient (PCC) of the two genes of *O. corymbosa* (*Olcov1051885* and *Olcov1029517*) versus all the other expressed genes.**

**Table S13. List of genes up- and down-regulated under cold 8C conditions.** The columns are: gene IDs, mean expression, log 2 fold change, standard error of log 2 fold change, p-value, adjusted p-value, direction, and function.

**Table S14. CETSA-MS analysis of fraction 34D against SKBR3 cell lysate.**

## Supplemental figures

**Figure S1.**
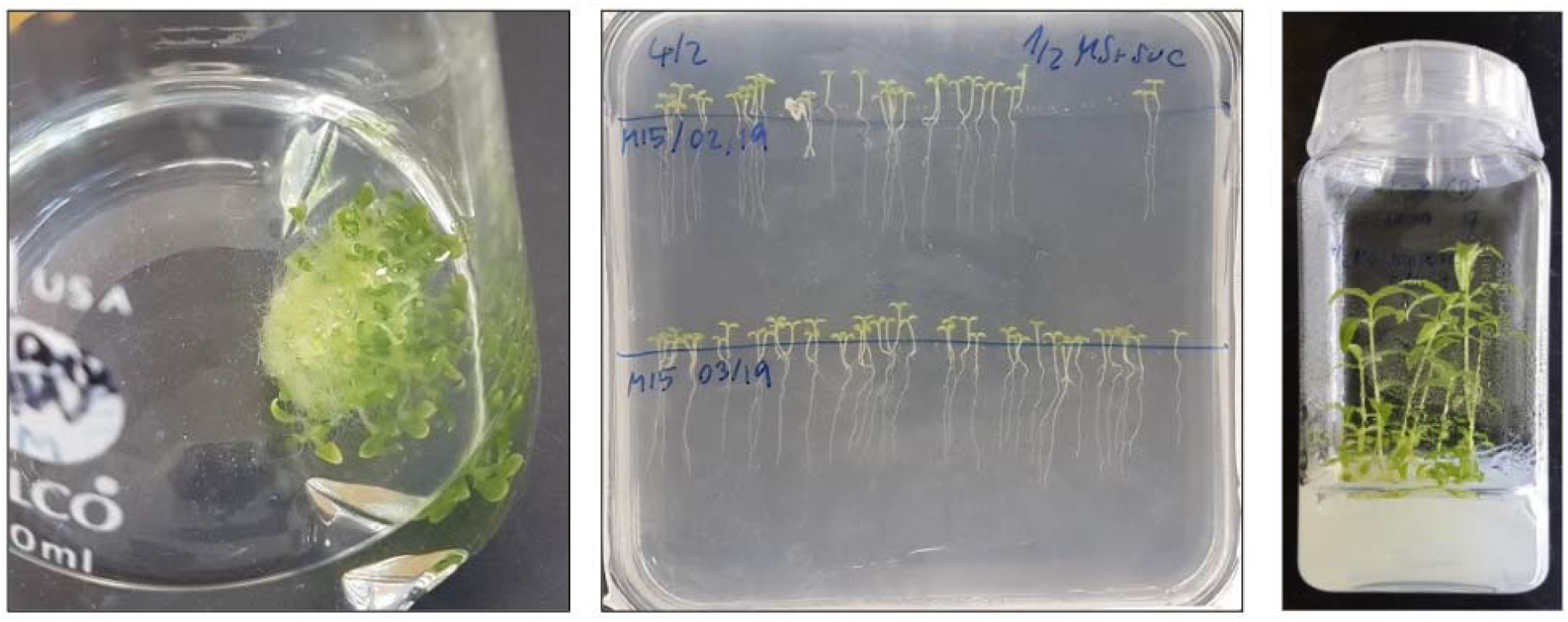
Liquid (left) and solid (middle, right) seedling growth conditions. For liquid culture about 100 seeds were used to inoculate 80 ml of ½ MS medium supplemented with 2.5 mM MES, 1% sucrose, and 1x MS vitamins (Sigma M3900), pH 5.8, and grown for 14 days at 100rpm, 28°C and 12h light (250 umol m^-2^ s^-1^). Seedlings were grown on plates or in jars on medium as for liquid cultures (supplemented with 0.8% agar) and grown for 10 days at 28°C and 12h light (250 umol m^-2^ s^-1^).

**Figure S2.**
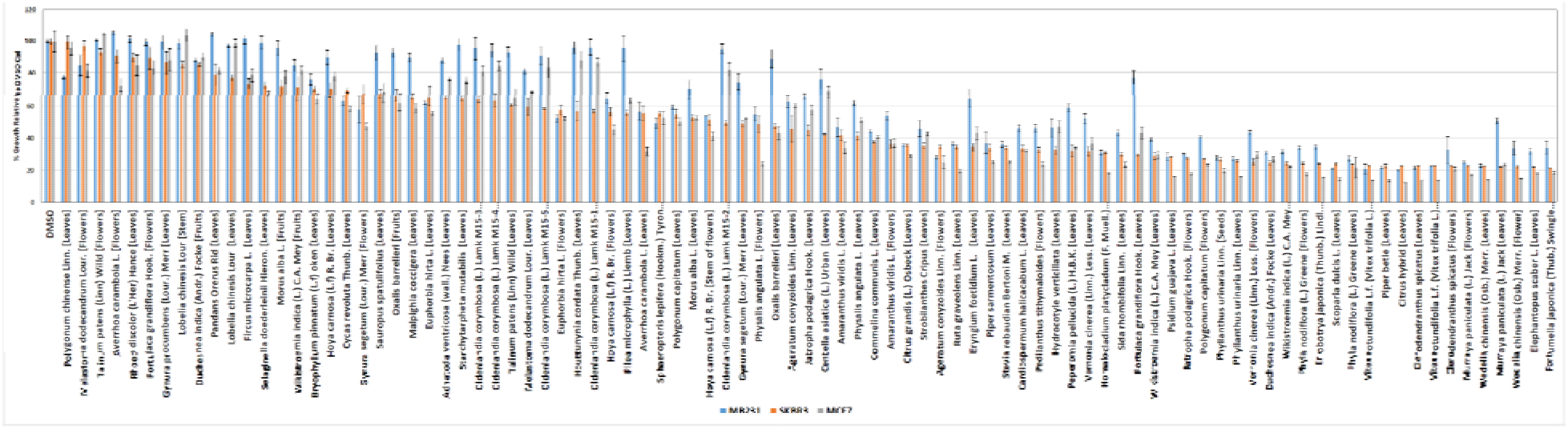
MTS assay comparison of *Oldenlandia corymbosa* to 61 medicinal plants using MB231, SKBR3, and MCF7 cancer cell lines. For each assay, at least three replicates were used. The error bars represent the standard deviation (*n*=3).

**Figure S3.**
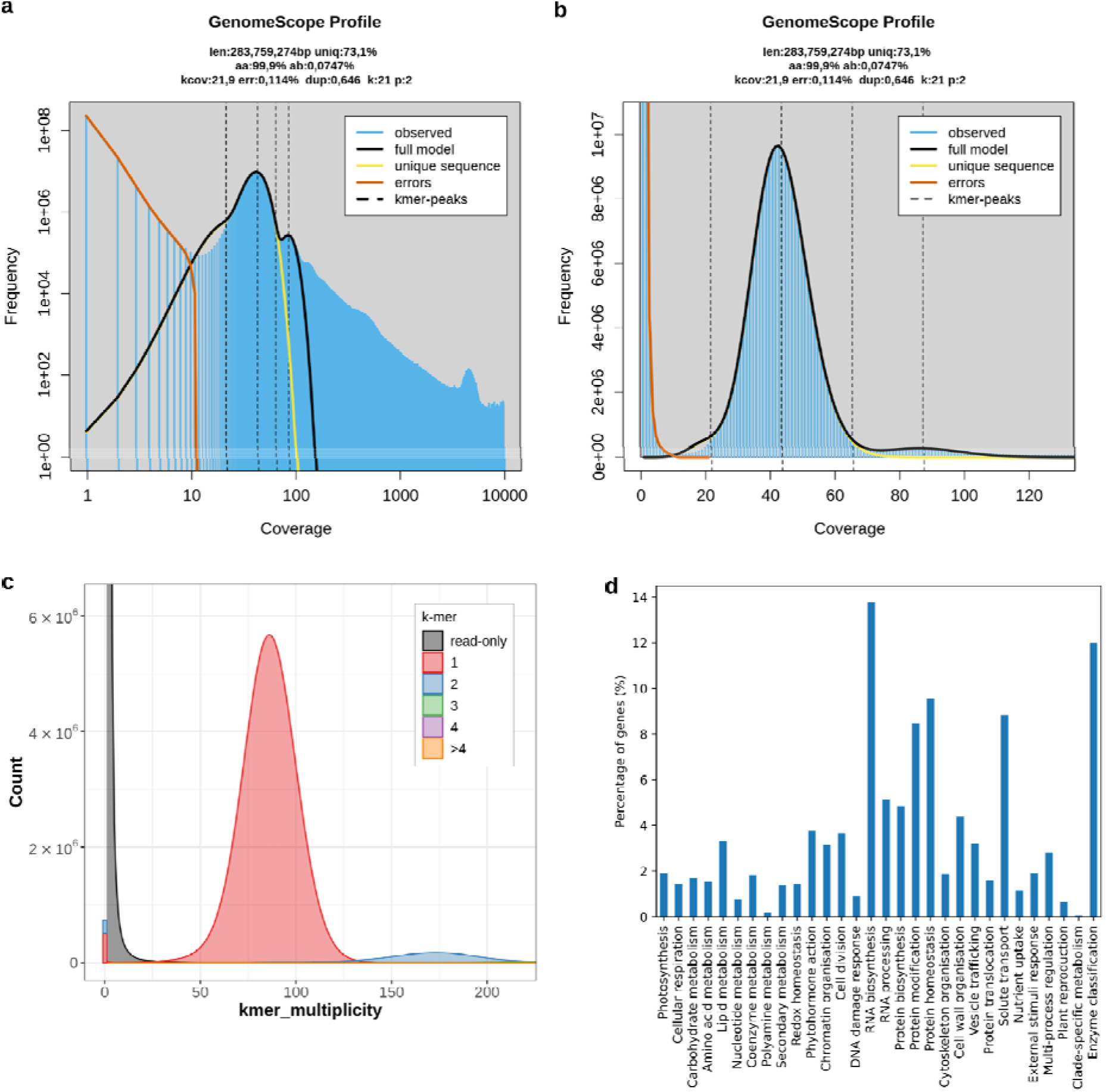
Genome analysis of *O. corymbosa*. a) GenomeScope *k*-mer profile plot using 21 mers showing the fit of the model (black) to the observed *k*-mer frequencies (blue). b) linear-scale plot focused on the main peaks. c) Merqury copy number spectrum (spectra-cn) plot of the genome assembly of O. corymbosa. Stacked histogram of the k-mer multiplicity collected from Illumina reads. The colors show the copy numbers found in the assembly. d) Percentage of genes annotated with MapMan bins. The percentages were calculated from the total number of genes with MapMan annotation (13,231).

**Figure S4.**
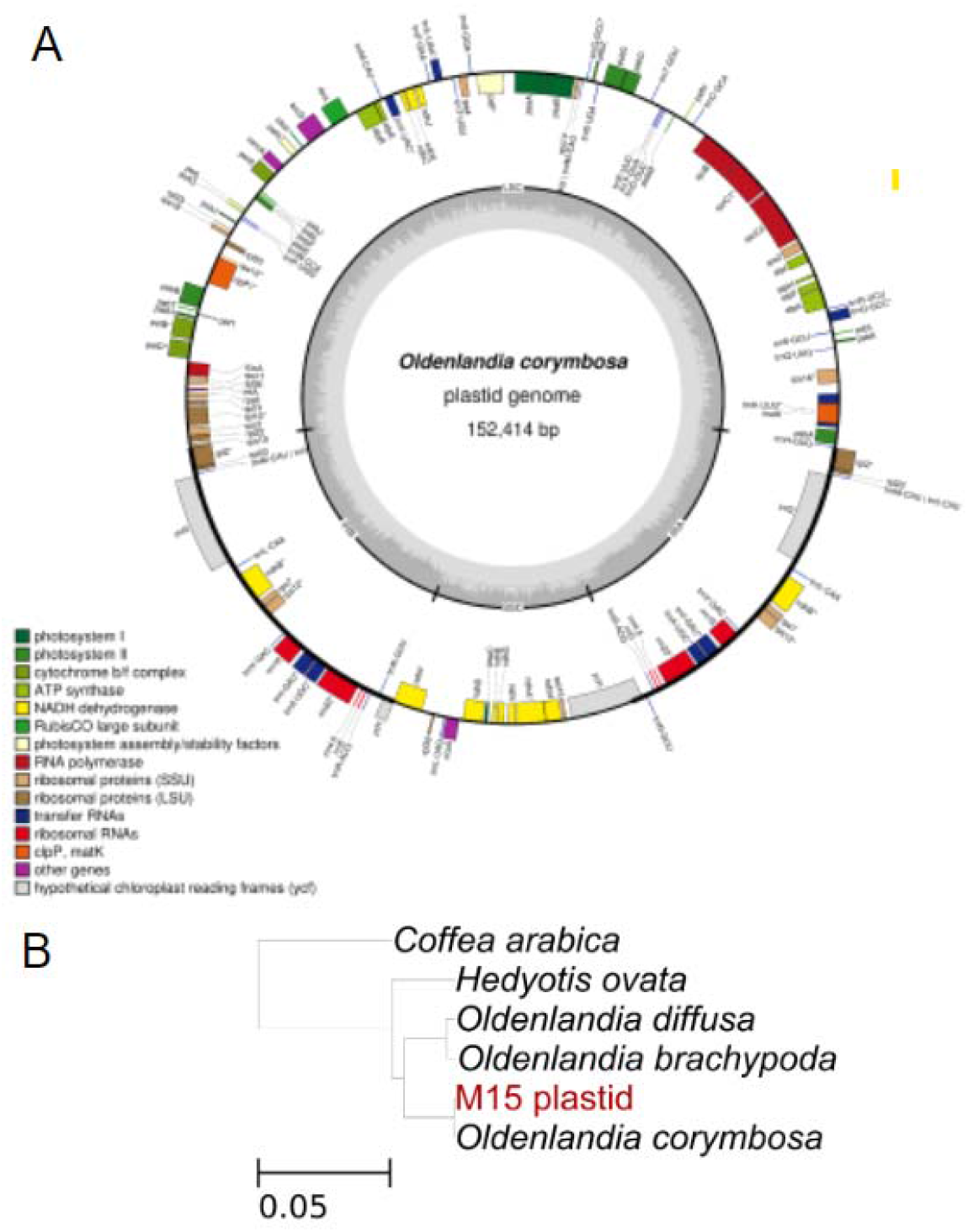
Analysis of plastid sequence of *Oldenlandia corymbosa*. A) Physical map of the plastid genome of *Oldenlandia corymbosa* as drawn by OGDRAW. The tick lines show the IR1 and IR2 regions, separating the SSC and LSC regions. Genes inside the circle are transcribed clockwise, while genes outside the circle are transcribed counterclockwise. Colours of the genes indicate their function. B) Species tree based on the plastid genome. All bootstrap values are maximal (100). The species sequenced and assembled in this project is marked in red (M15 plastid).

**Figure S5.**
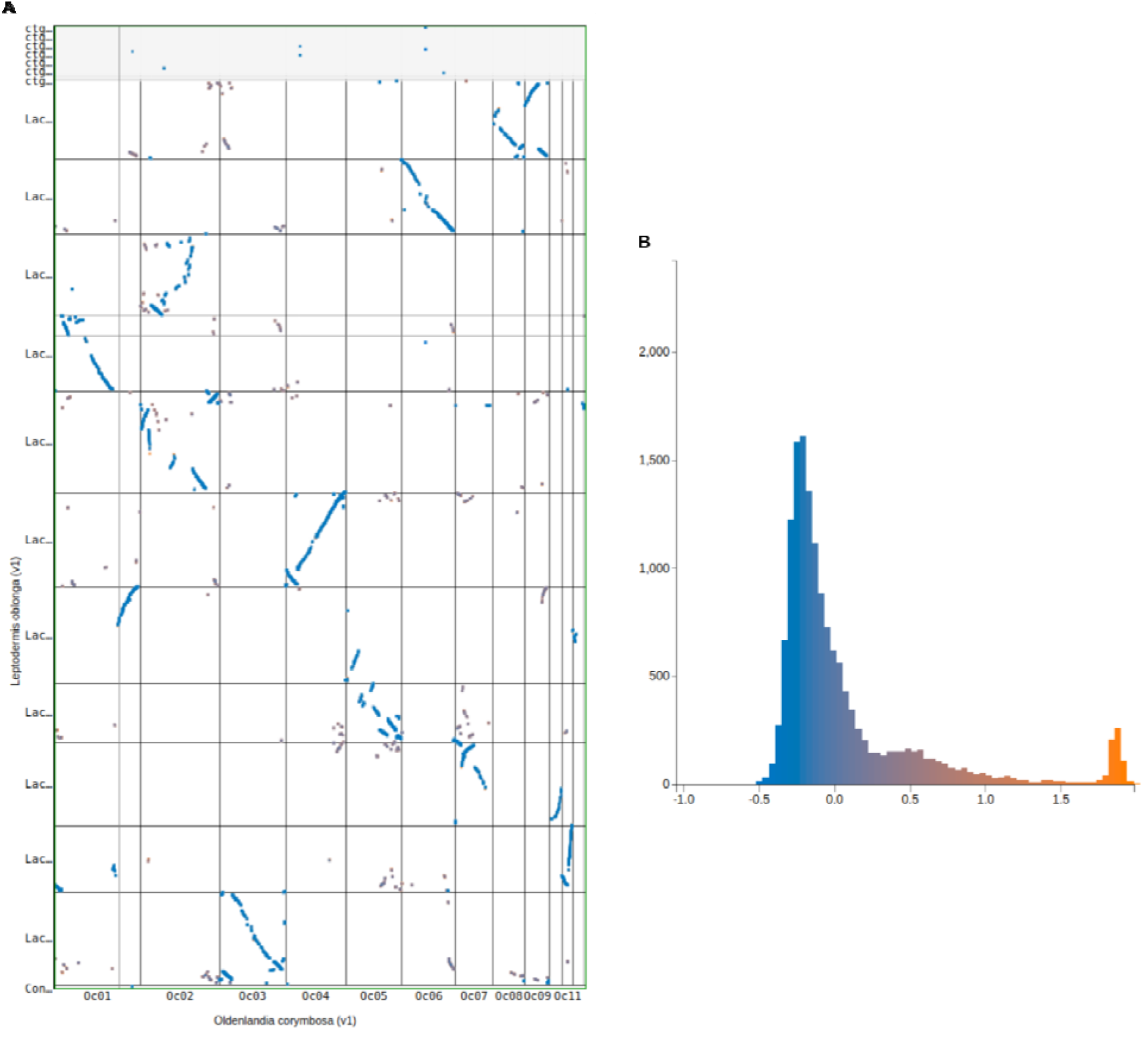
Synteny and Ks analysis between *O. corymbosa* and *L. oblonga*. A. Syntenic dotplot with Ks coloration between *O. corymbosa* (x-axis) and *Leptodermis oblonga* (y-axis). Blue lines show orthologous genes between both species. B. Histogram of synonymous mutation rates (Ks) for syntenic genes between *O. corymbosa* and *L. oblonga*.

**Figure S6.**
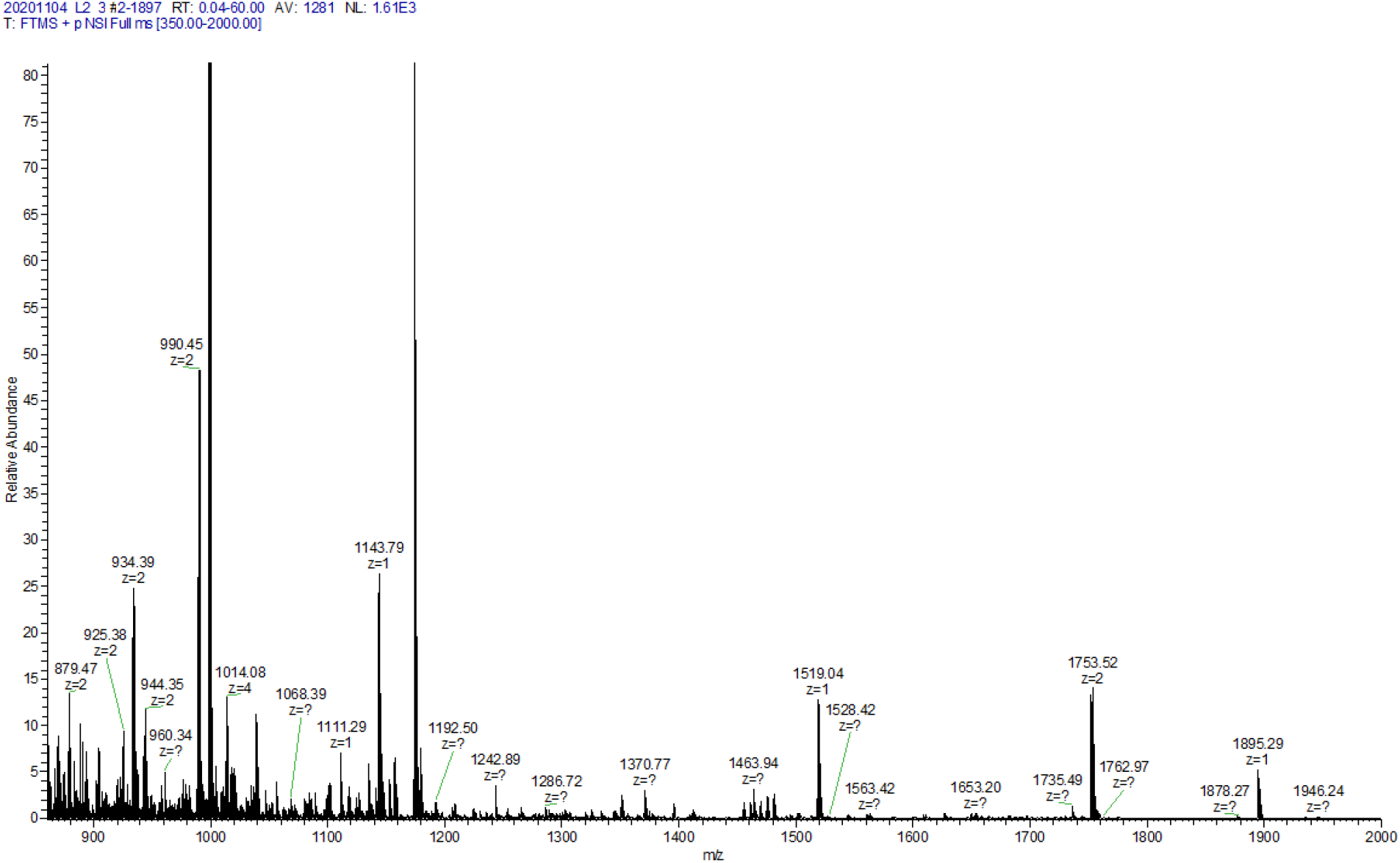
LC-MS/MS spectra of *Oldenlandia corymbosa* leaf samples identifies a potential cyclotide. The 3507 Da cyclotide is represented by the 1753.52 (z = 2) peak.

**Figure S7.**
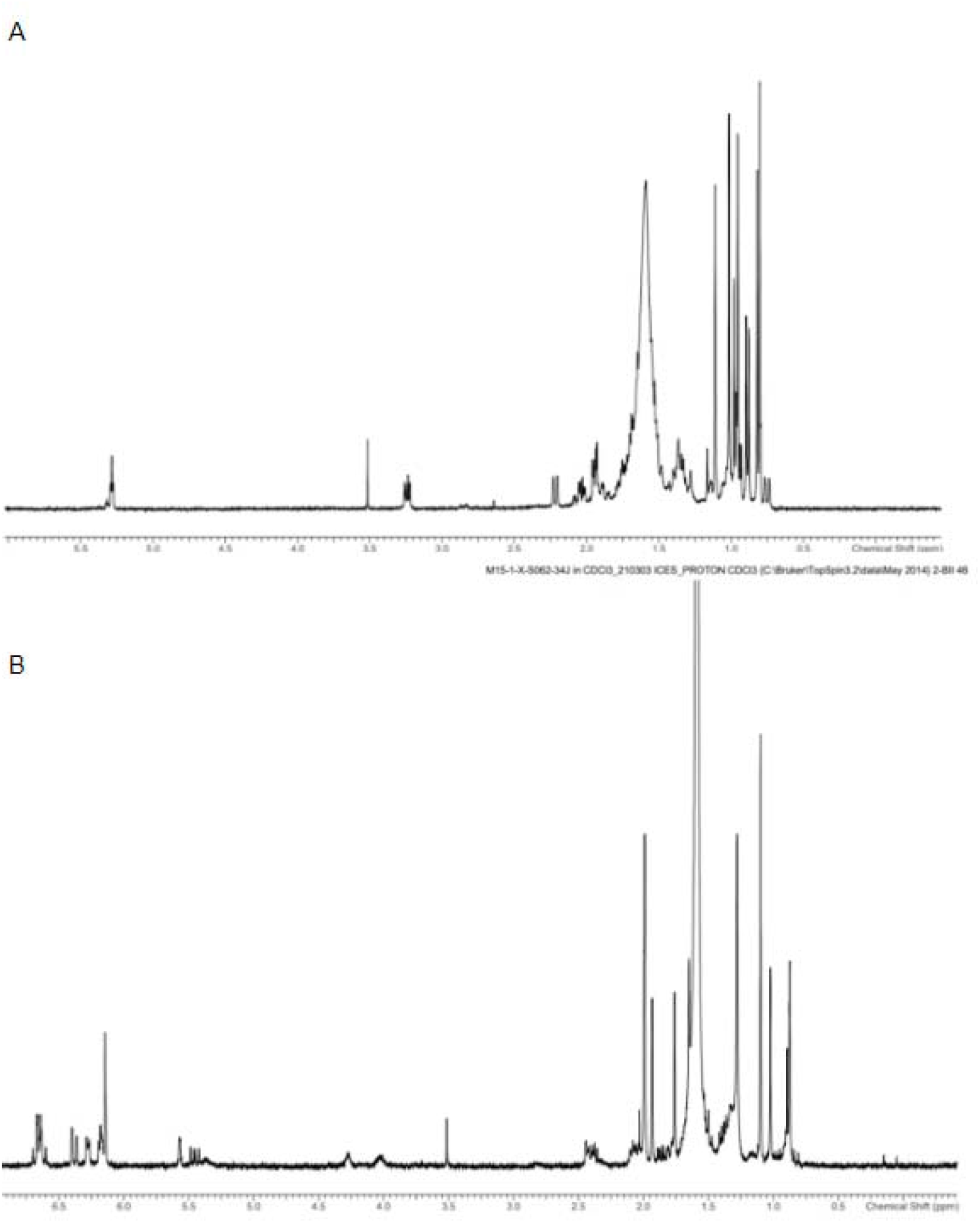
1H NMR spectra of O. corymbosa fractions. A) Fraction 34D ursolic acid in CDCl3. B) Fraction 34J lutein in CDCl3.

**Figure S8.**
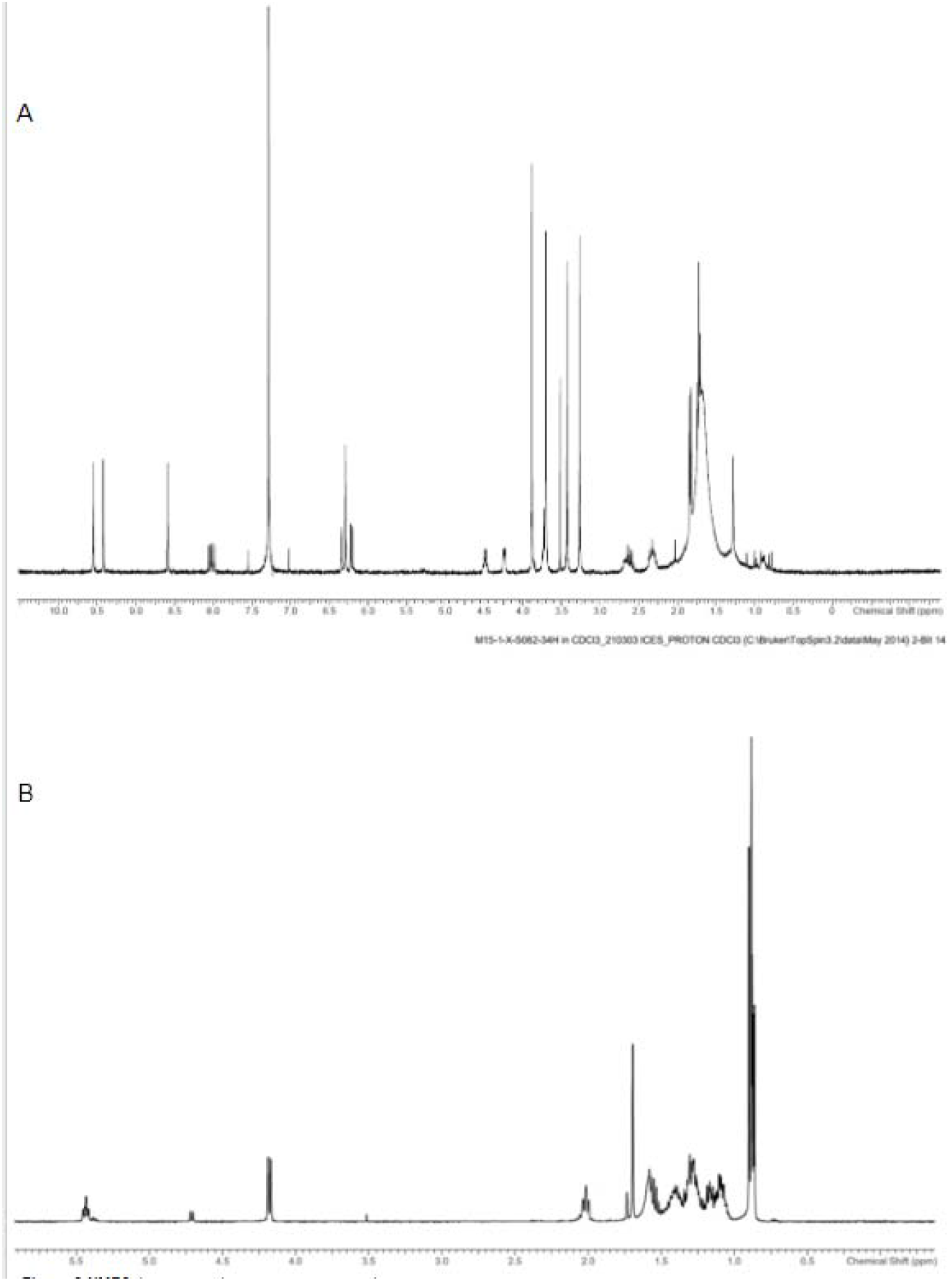
1H NMR spectra of O. corymbosa fractions. A) Fraction 34F (Pheophorbide L) in CDCl3. B) 34H (Phytol) in CDCl3

**Figure S9.**
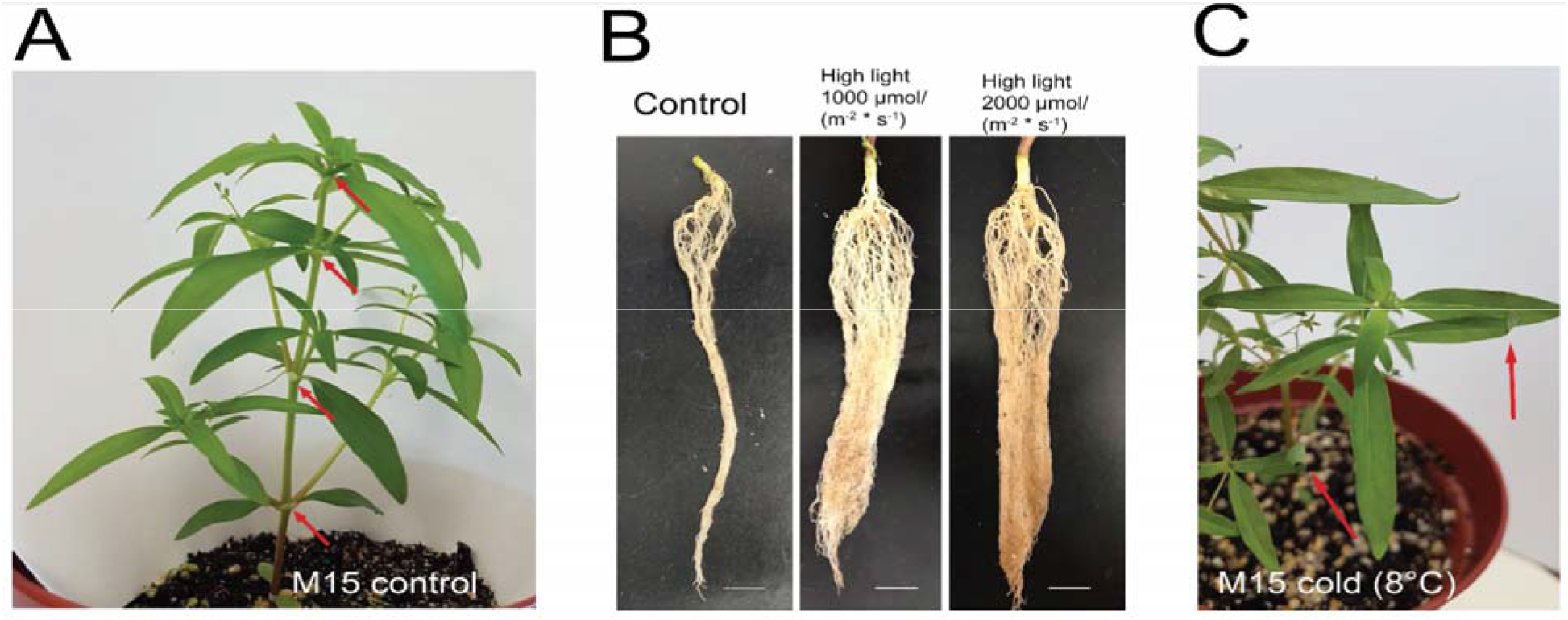
*O. corymbosa* phenotypes caused by abiotic stress. A) Plant with 4th node developed at the start of the stress experiments. B) Comparison of roots from control, and two high light conditions. C) Rolled-in leaves (indicated by red arrows) of a plant treated at 8°C.

**Figure S10.**
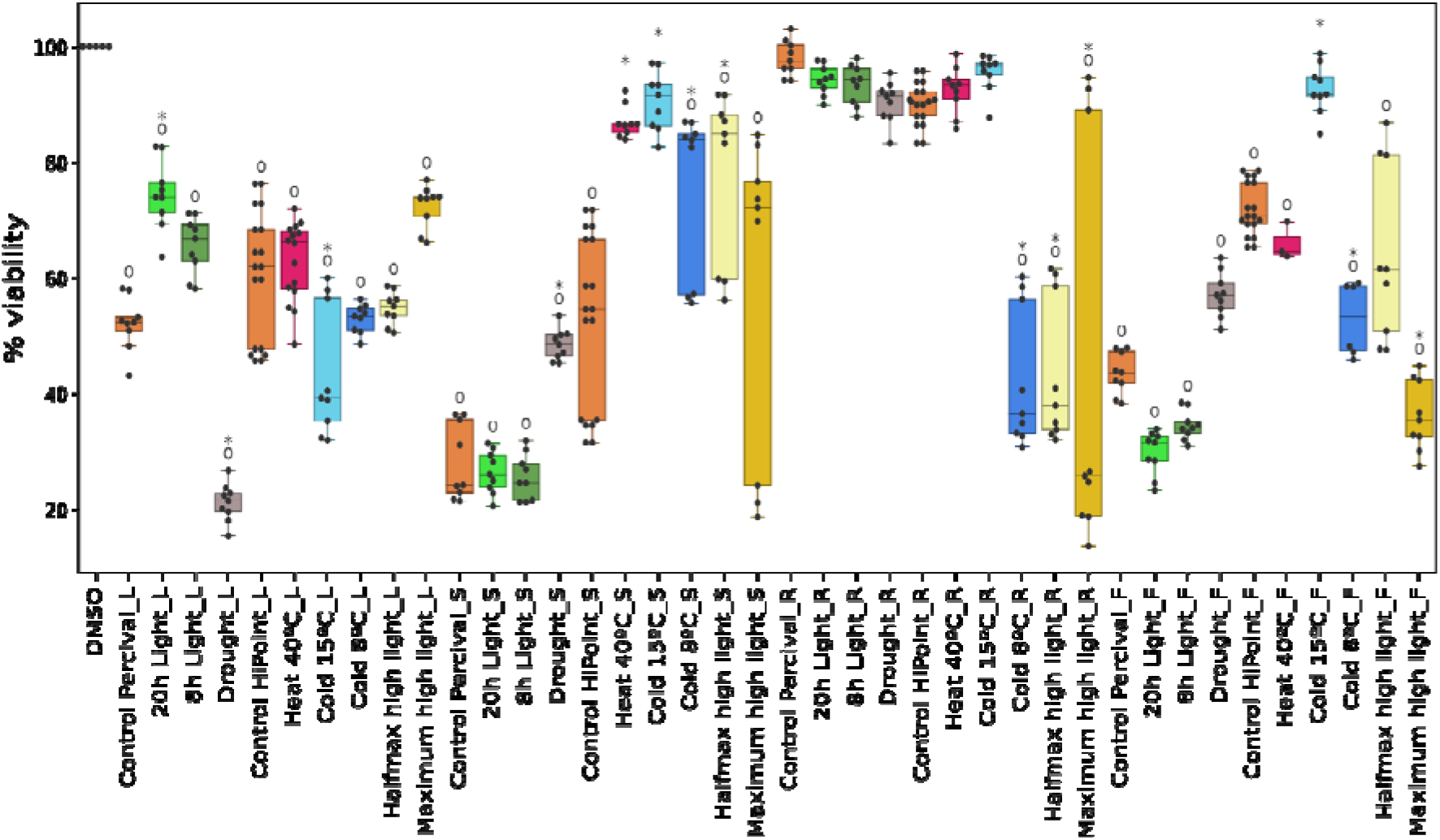
MTT assay activities against SKBR3 cancer cell lines. Samples are indicated on the x-axis, while cell viability is shown on the y-axis. The black dots indicate one measurement, while the boxplots summarize the distribution of the data. For each sample, three plants were collected, and the activity of each sampled organ was measured with three MTT assay measurements, giving nine measurements per treatment and organ. NSI_L indicates leave extracts from Controls in Percival; 20h_L, leaves from 20L/4D; 8h_L, leaves from 8L/16D; Dr_L, leaves from drought; NSII_L, leaves from Controls in HiPoint chamber; H40_L, leaves from heat stress; C15_L, leaves from cold at 15°C; C8_L, leaves cold at 8°C; hHL_L, leaves from half-maximum high light (1000 μmol·m^-2^·s^-1^); mHL_L, leaves from maximum high light (2000 μmol·m^-2^·s^-1^). S, stems; F, flowers; R, roots. Circles and asterisks indicate significant differences (p<0.05) between the treatment and the DMSO control, and between plant extract control and treatment, respectively.

**Figure S11.**
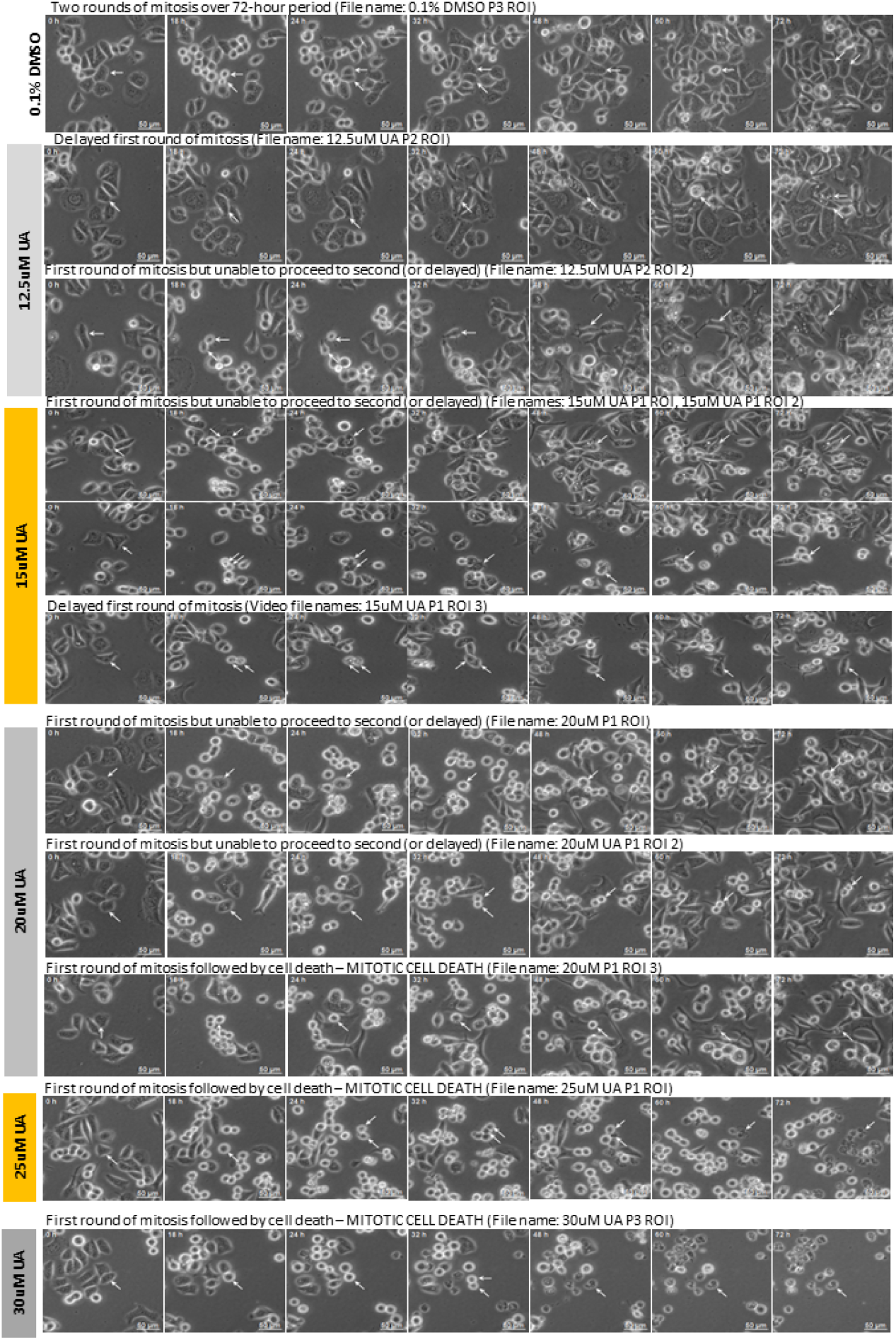
Phase contrast images of SKBR3 cells treated with ursolic acid and DMSO control over 72 hours. Rows represent different concentrations, while columns correspond to timepoints. The white arrows indicate the discussed cells.

**Figure S12.**
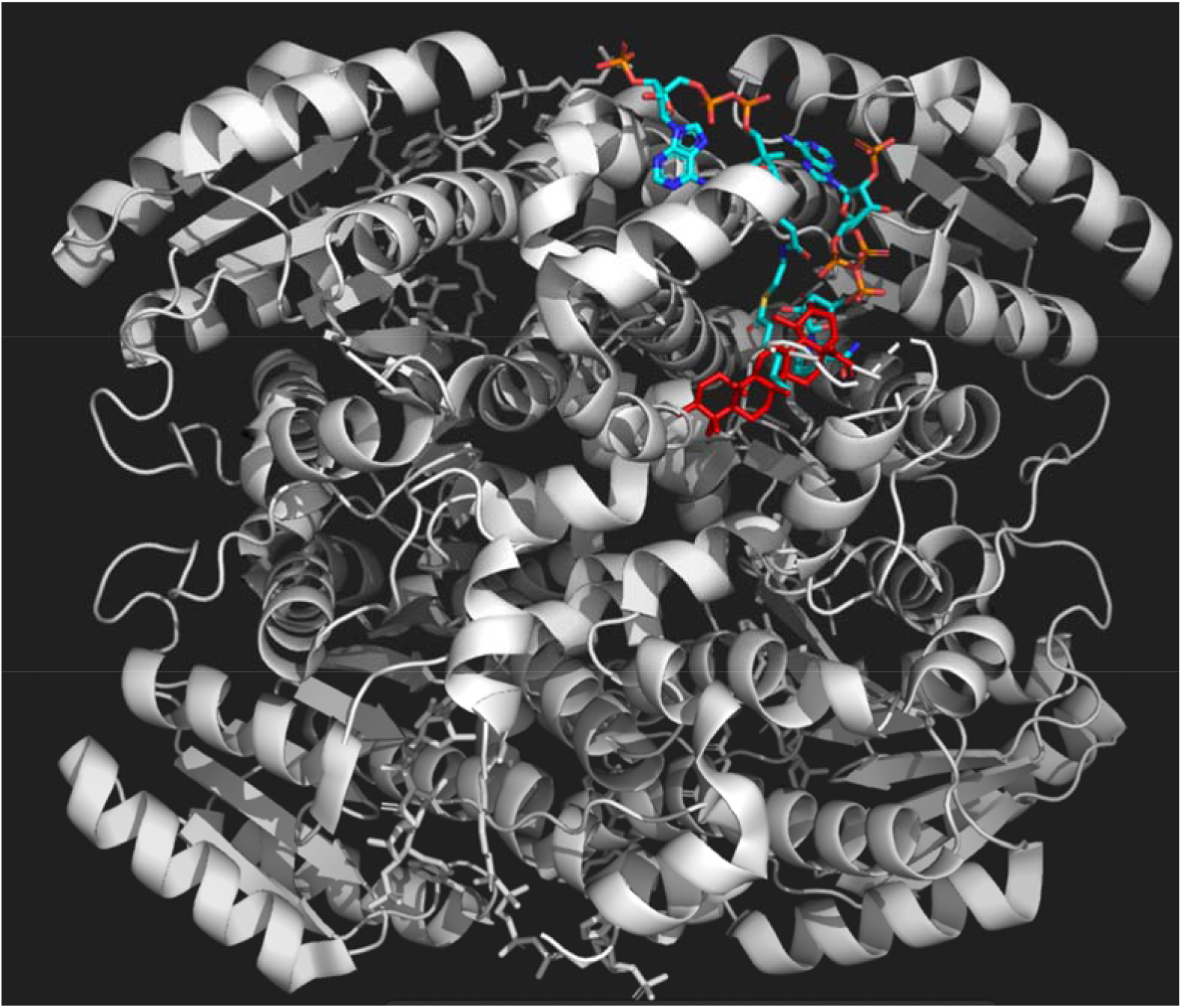
**Atomic model of DECR1 in complex with natural ligand and ursolic acid**. The protein is colored white, ursolic acid is colored red, while the natural ligands are colored orange (NADP nicotiamide-adenine-dinucleotide phosphate) and cyan (hexanoyl-coenzyme A).

## Supplemental data

**Supplemental Data 1.** Video 1. Phase-contrast movie of SKBR3 cells treated with 0.1% DMSO over 72 hours.

**Supplemental Data 2.** Video 2. Phase-contrast movie of SKBR3 cells treated with 0.1% DMSO over 72 hours.

**Supplemental Data 3.** Pymol session file containing the structures of DECR1, Maspardin and HIBCH bound by ursolic acid.

