## Supplementary material for "Genomic, transcriptomic, and metabolomic analysis of Traditional Chinese Medicine plant *Oldenlandia corymbosa* reveals the biosynthesis and mode of action of anti-cancer metabolites": Figure S1-12

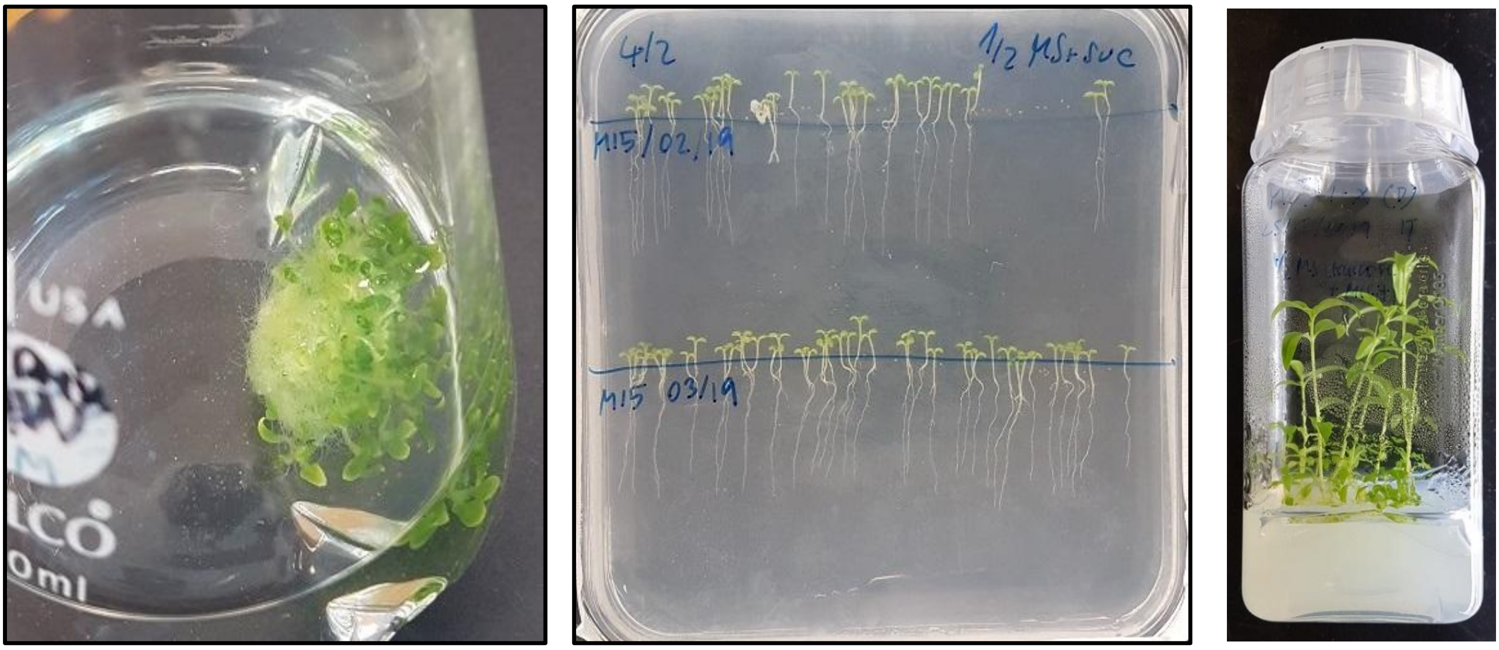


**Figure S1. Liquid (left) and solid (middle, right) seedling growth conditions.** For liquid culture about 100 seeds were used to inoculate 80 ml of ½ MS medium supplemented with 2.5 mM MES, 1% sucrose, and 1x MS vitamins (Sigma M3900), pH 5.8, and grown for 14 days at 100rpm, 28°C and 12h light (250 umol $m^{-2}\,s^{-1}$). Seedlings were grown on plates or in jars on medium as for liquid cultures (supplemented with 0.8% agar) and grown for 10 days at 28°C and 12h light (250 umol $m^{-2}\,s^{-1}$).

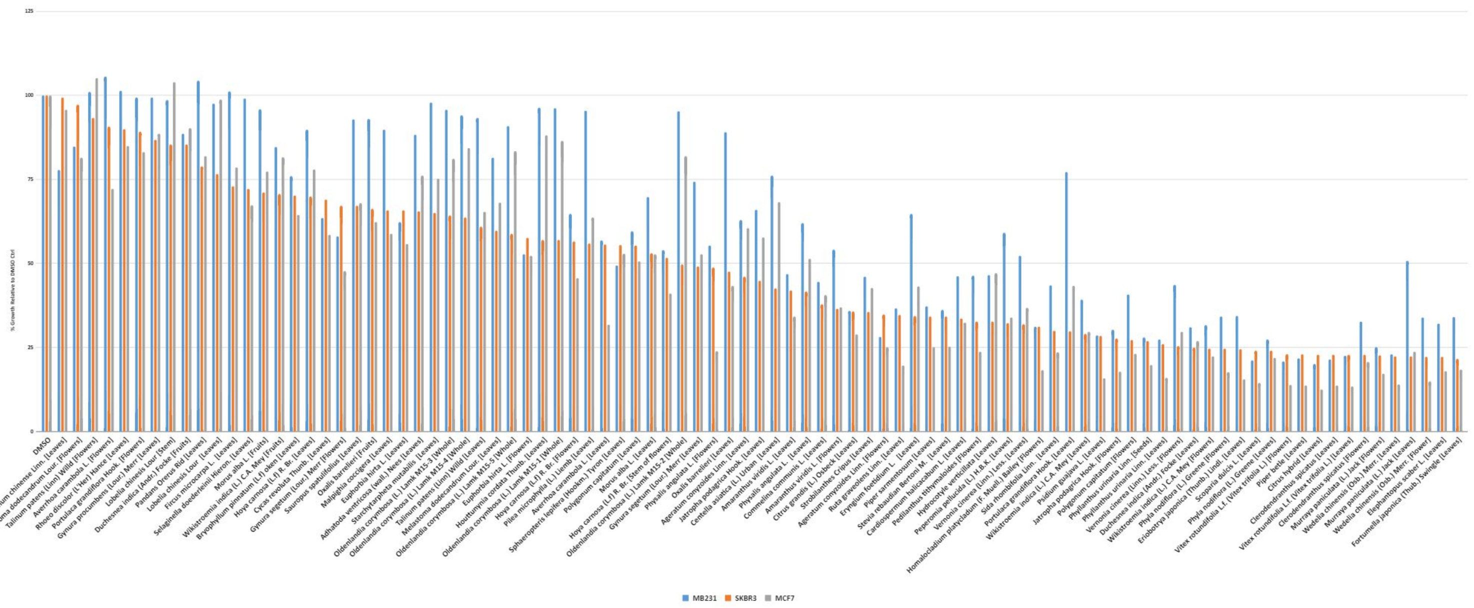


**Figure S2. MTS assay comparison of *Oldenlandia corymbosa* to 61 medicinal plants using MB231, SKBR3, and MCF7 cancer cell lines.** For each assay, at least three replicates were used. The error bars represent standard deviation (*n*=3).

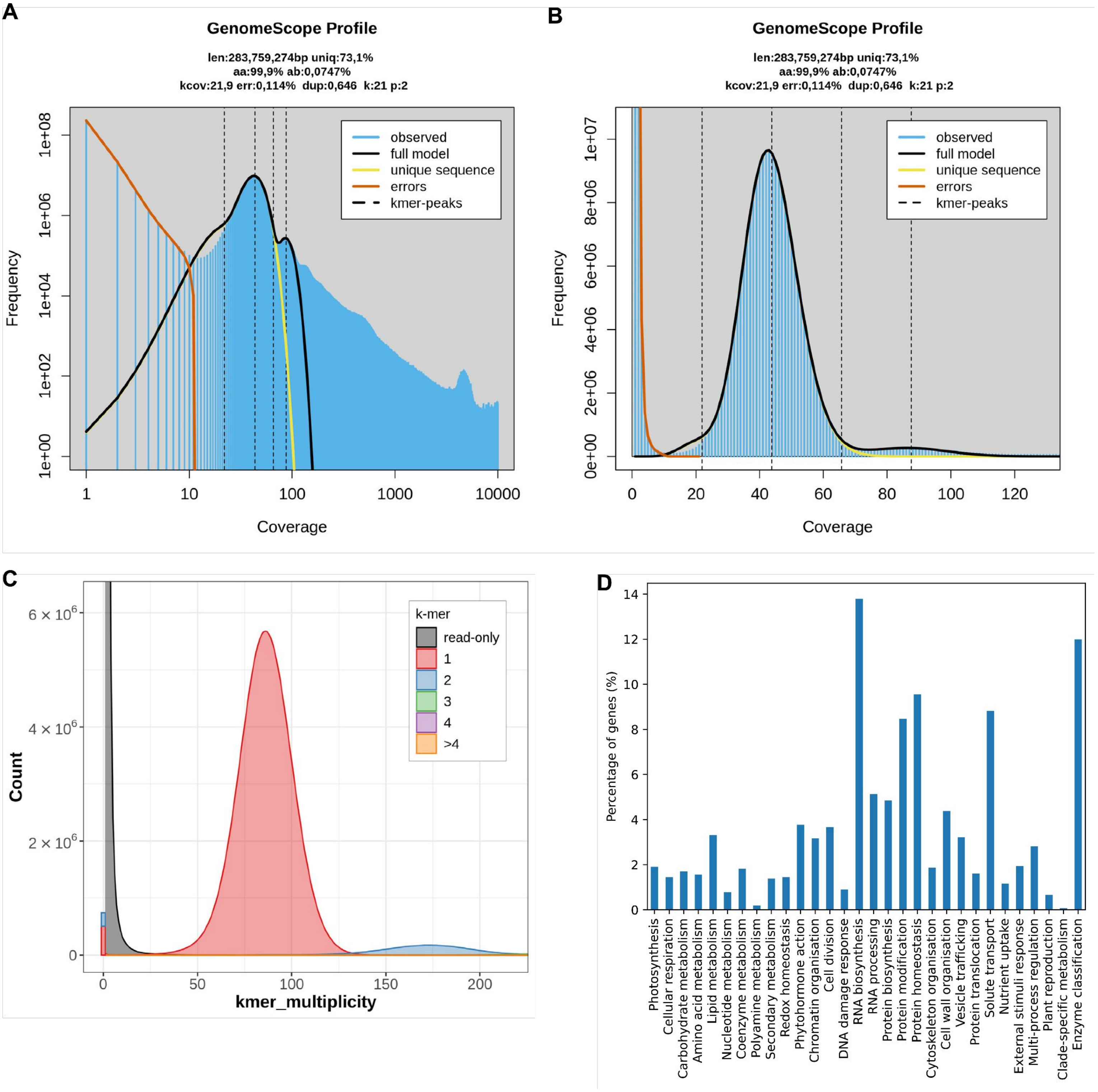


**Figure S3. Genome analysis of *O. corymbosa*.** A) GenomeScope *k*-mer profile plot using 21 mers showing the fit of the model (black) to the observed *k*-mer frequencies (blue). B) linear-scale plot focused on the main peaks. C) Merqury copy number spectrum (spectra-cn) plot of the genome assembly of O. corymbosa. Stacked histogram of the k-mer multiplicity collected from Illumina reads. The colors show the copy numbers found in the assembly. D) Percentage of genes annotated with MapMan bins. The percentages were calculated from the total number of genes with MapMan annotation (13,231).

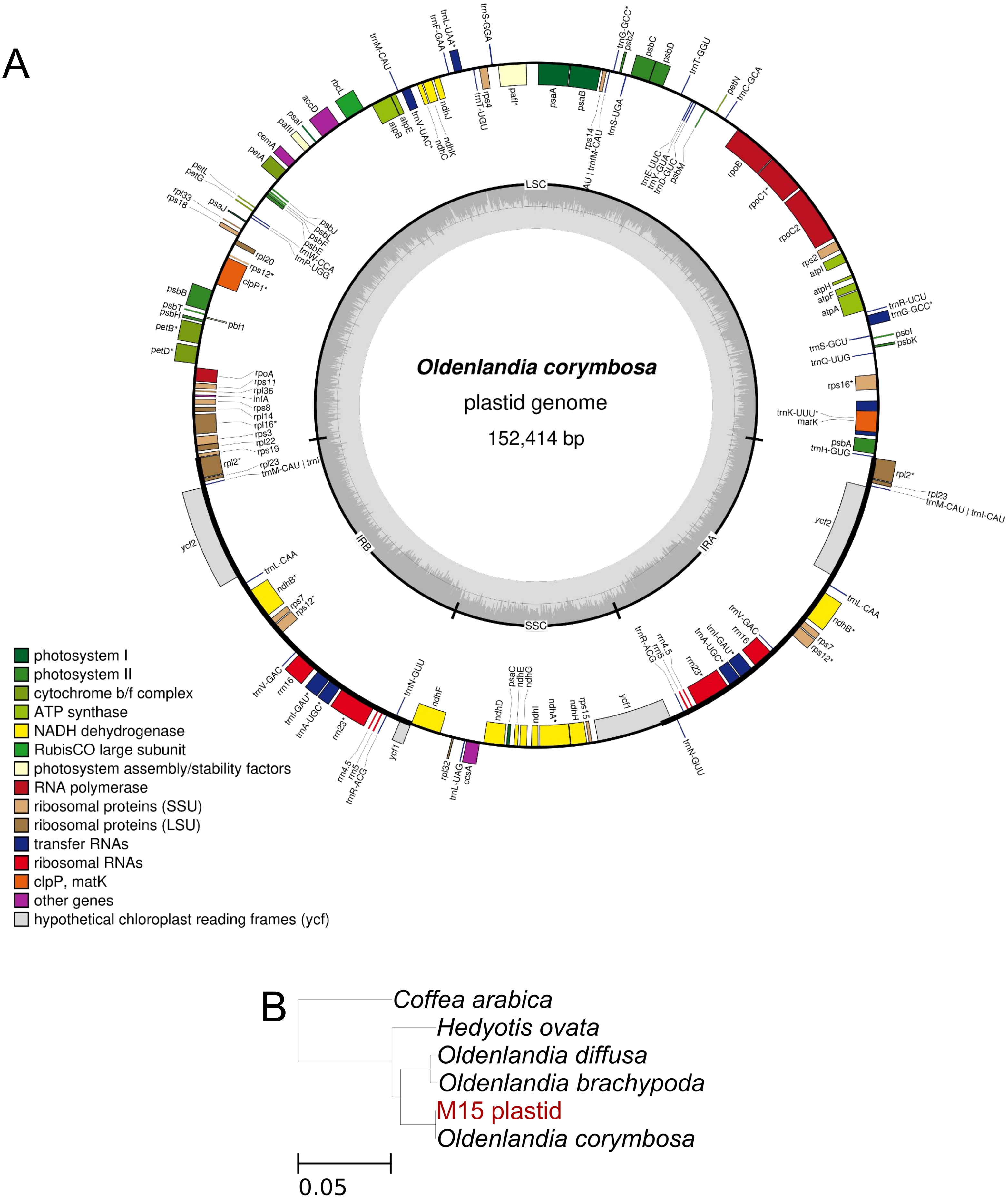


**Figure S4. Analysis of plastid sequence of *Oldenlandia corymbosa*.** A) Physical map of the plastid genome of *Oldenlandia corymbosa* as drawn by OGDRAW. The tick lines show the IR1 and IR2 regions, separating the SSC and LSC regions. Genes inside the circle are transcribed clockwise, while genes outside the circle are transcribed counterclockwise. Colours of the genes indicate their function. B) Species tree based on the plastid genome. All bootstrap values are maximal (100). The species sequenced and assembled in this project is marked in red (M15 plastid).

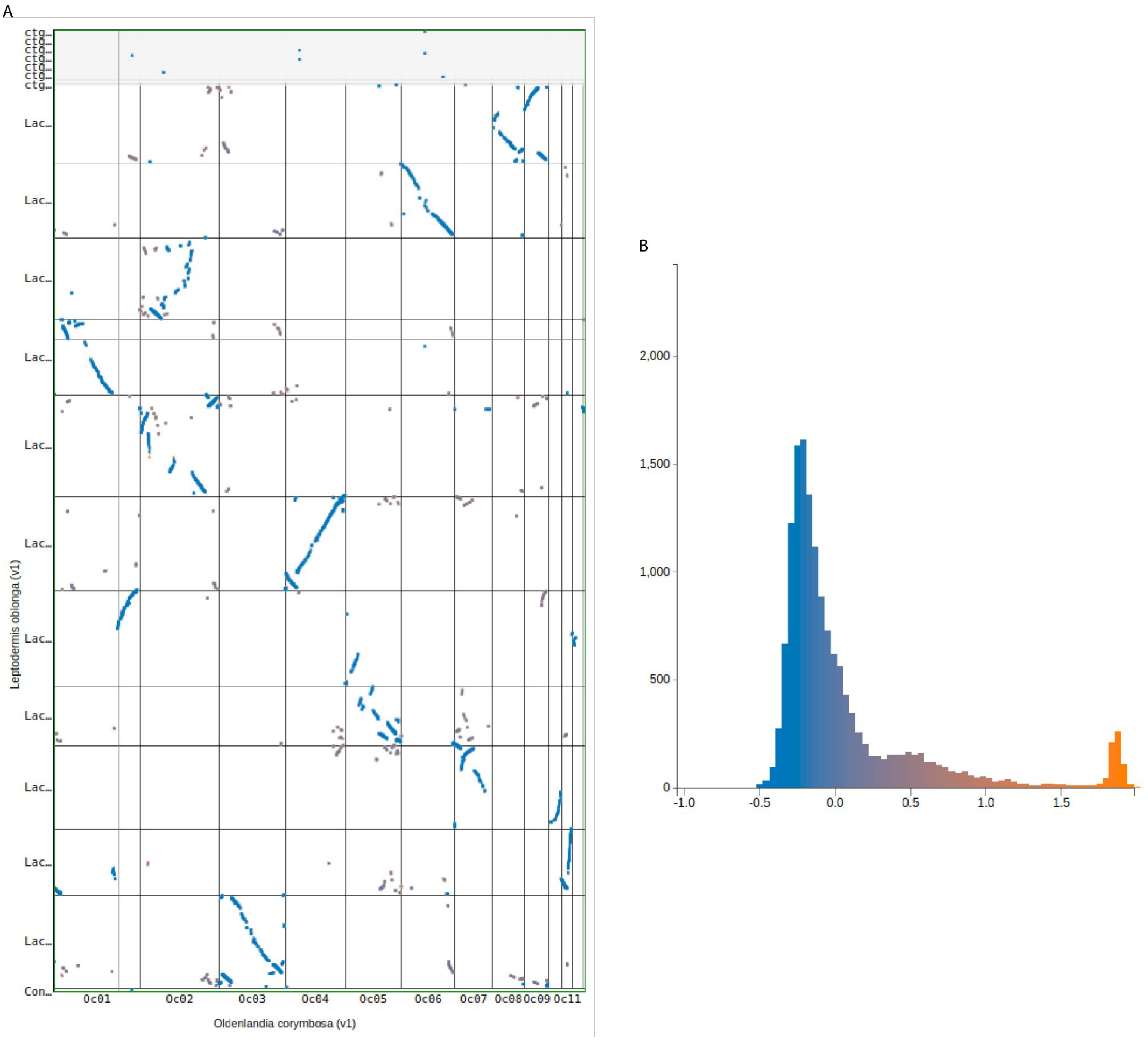


**Figure S5. Synteny and Ks analysis between *O. corymbosa* and *L. oblonga*.** A. Syntenic dotplot with Ks coloration between *O. corymbosa* (x-axis) and *Leptodermis oblonga* (y-axis). Blue lines show orthologous genes between both species. B. Histogram of synonymous mutation rates (Ks) for syntenic genes between *O. corymbosa* and *L. oblonga*.

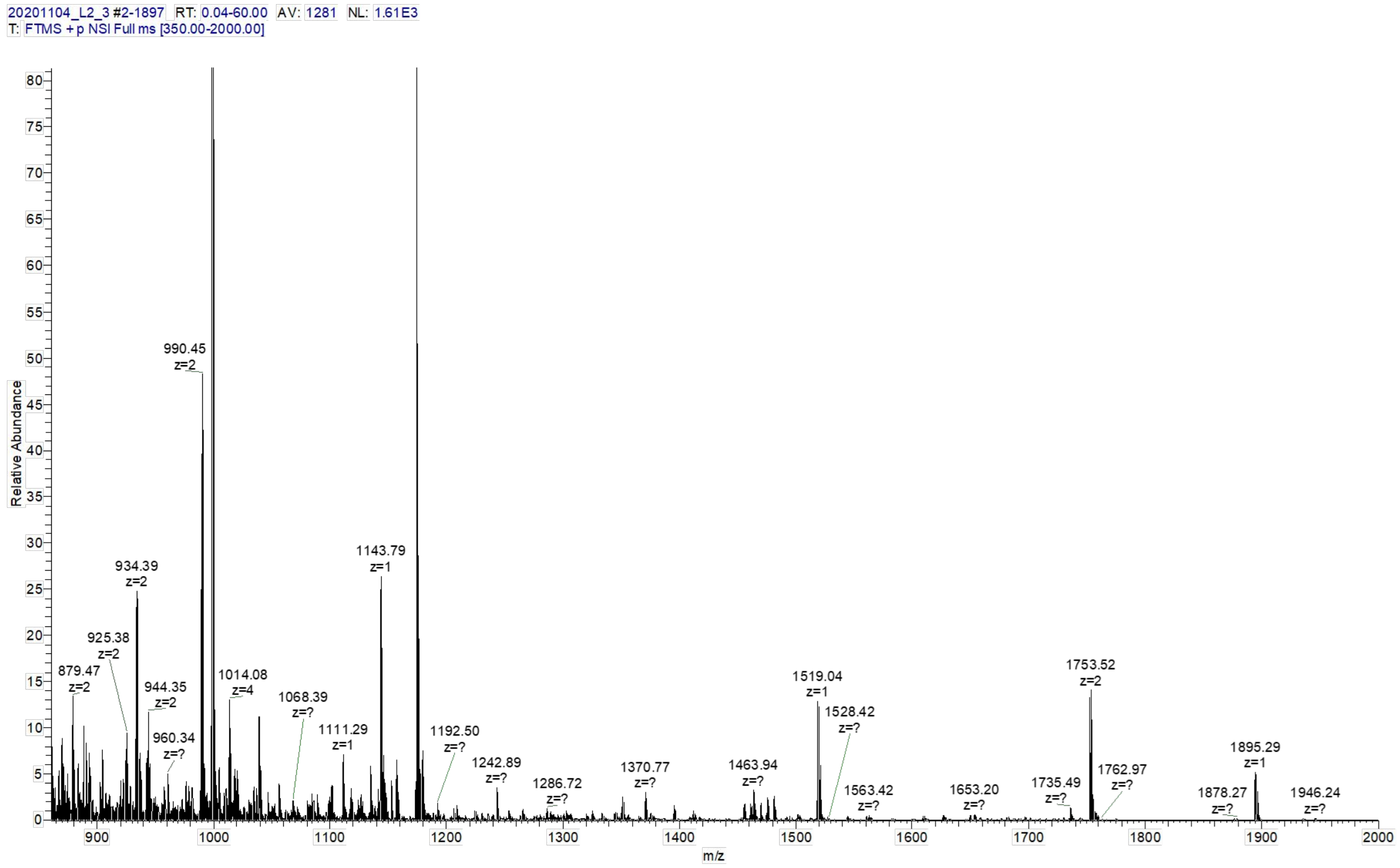


**Figure S6. LC-MS/MS spectra of *Oldenlandia corymbosa* leaf samples identifies a potential cyclotide.** The 3507 Da cyclotide is represented by the 1753.52 (z = 2) peak.

A

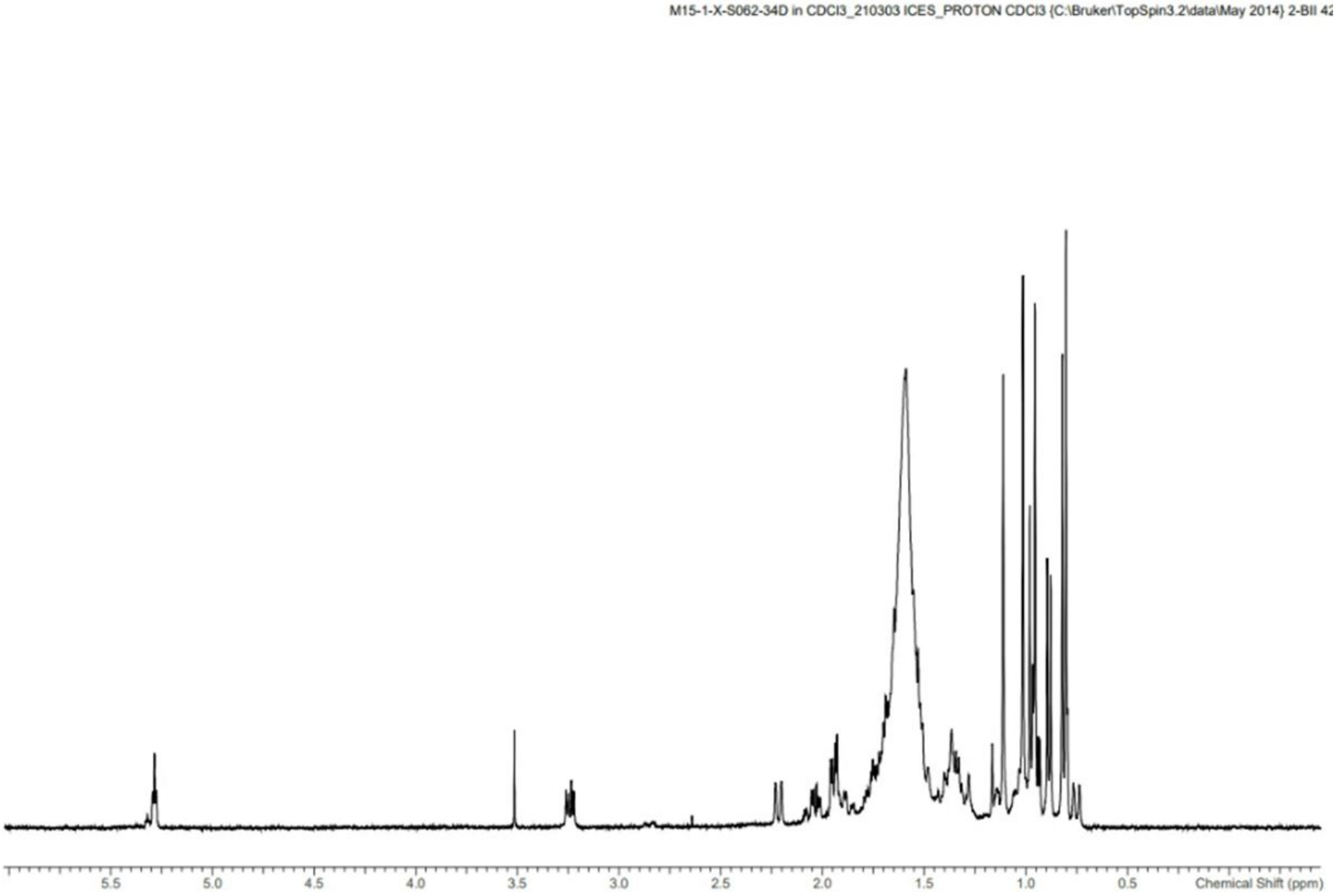


B

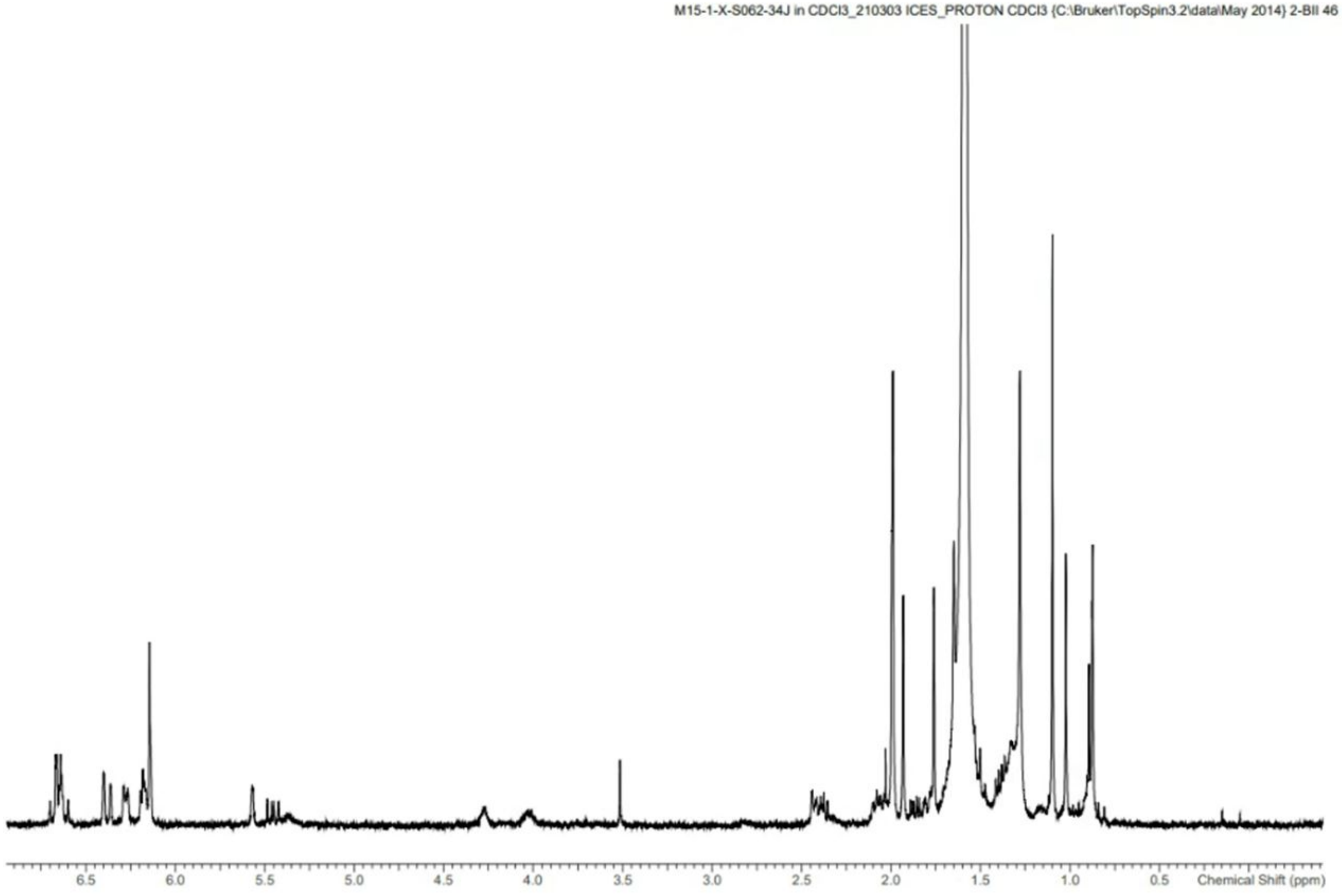


**Figure S7. 1H NMR spectra of O. corymbosa fractions.** A) Fraction 34D ursolic acid in CDCl3. B) Fraction 34J lutein in CDCl3.

A

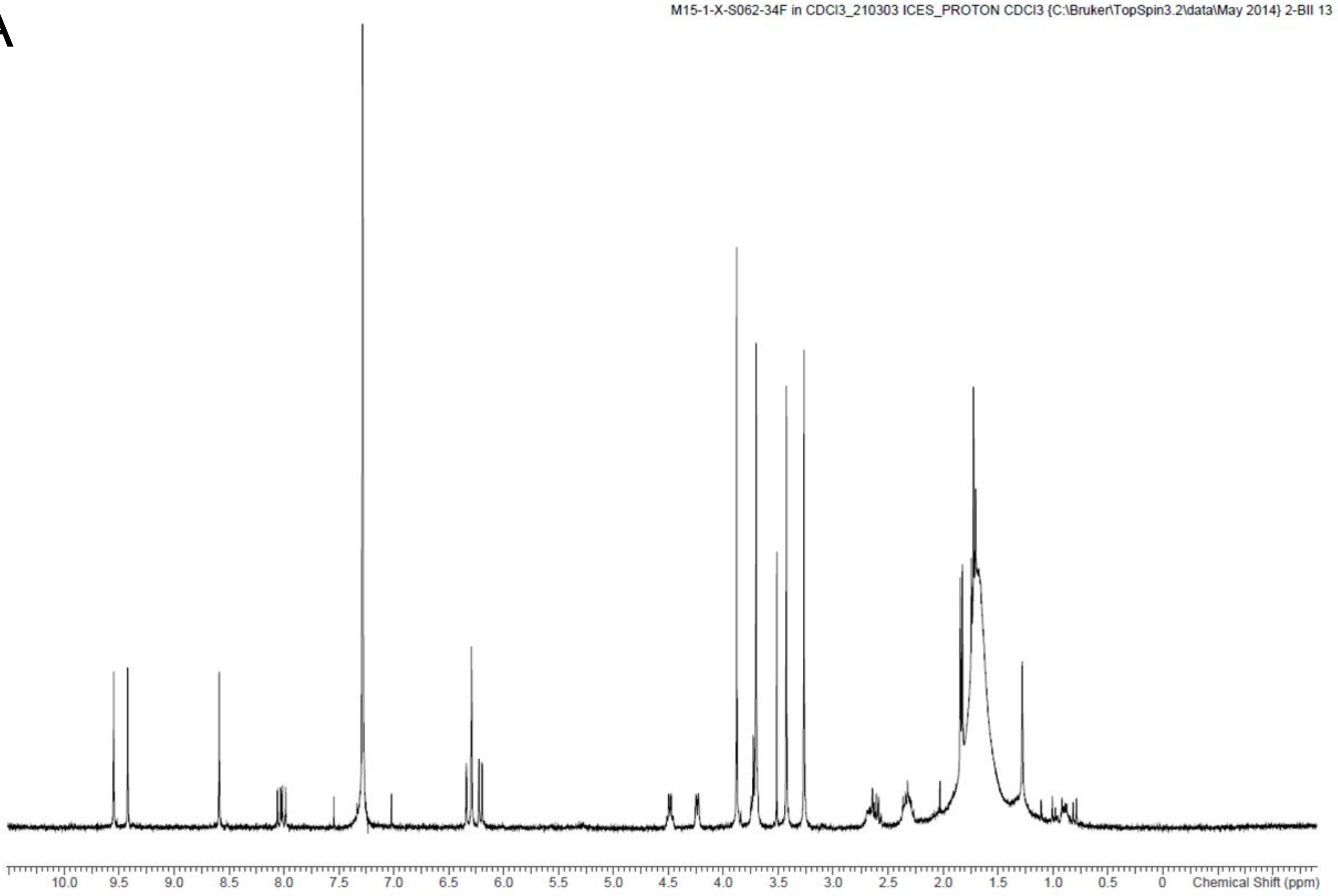


B

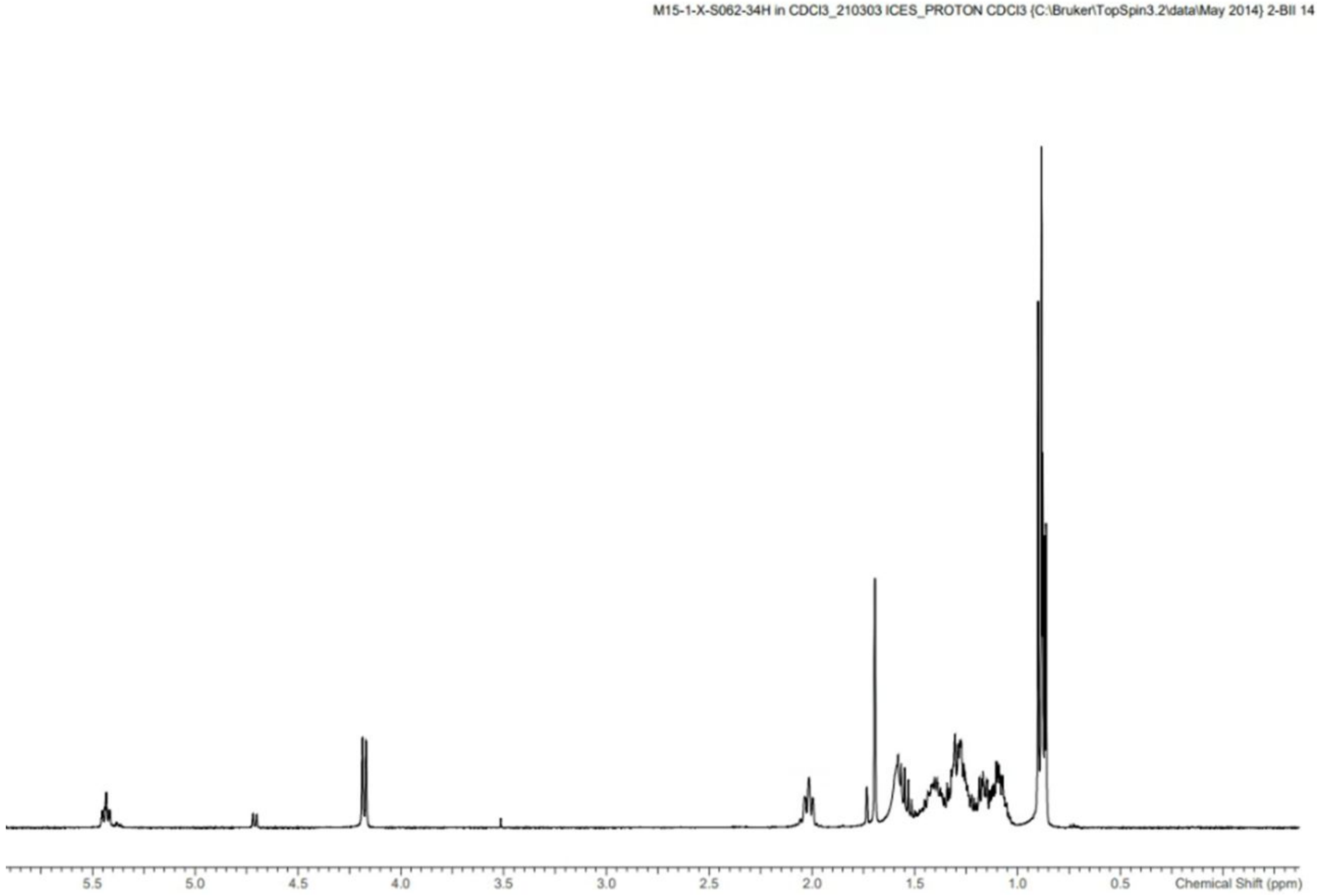


**Figure S8. 1H NMR spectra of O. corymbosa fractions.** A) Fraction 34F (Pheophorbide α) in CDCl3. B) 34H (Phytol) in CDCl3

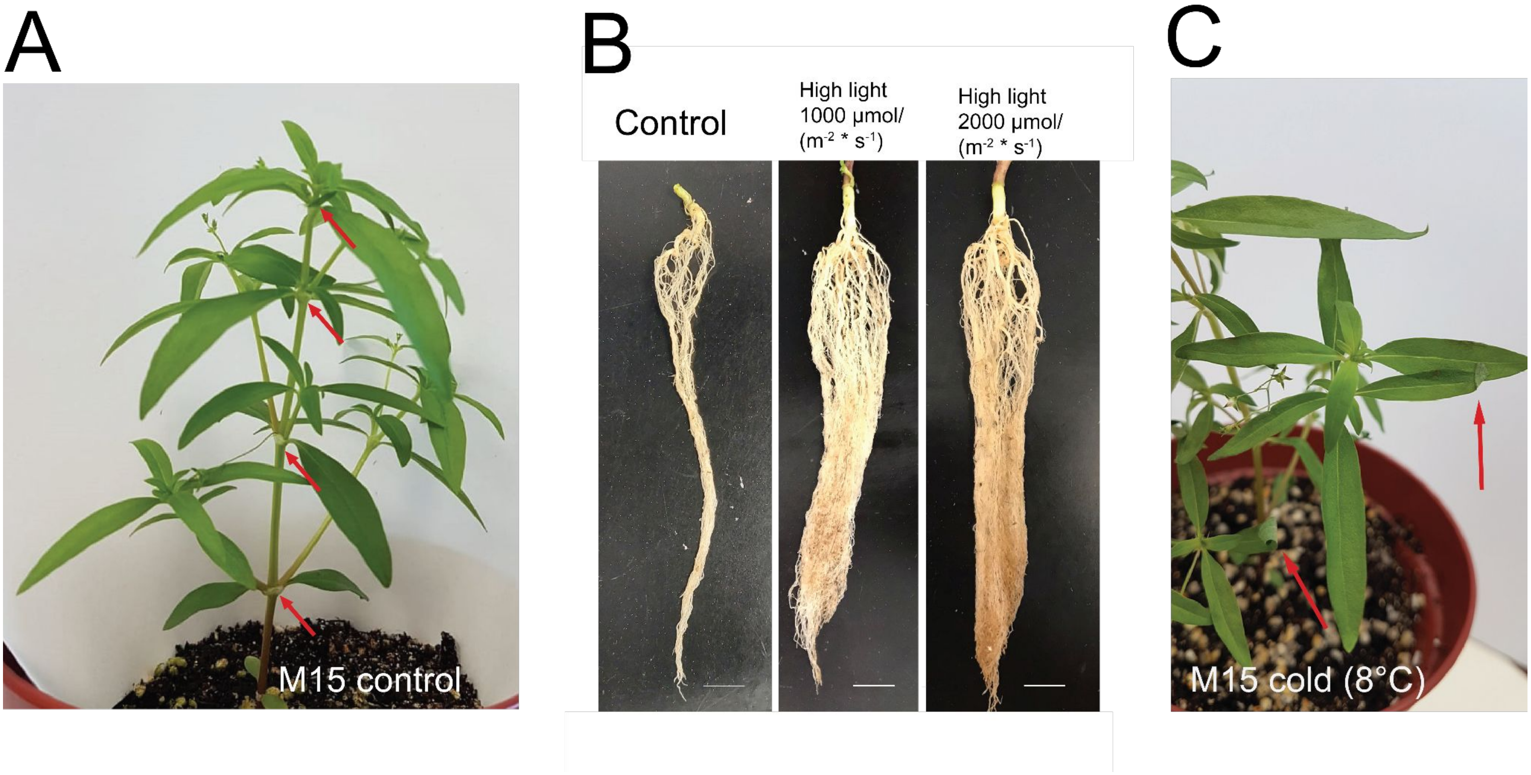


**Figure S9.** ***O. corymbosa*** **phenotypes caused by abiotic stress.** A) Plant with 4th node developed at the start of the stress experiments. B) Comparison of roots from control, and two high light conditions. C) Rolled-in leaves (indicated by red arrows) of plant treated at 8°C.

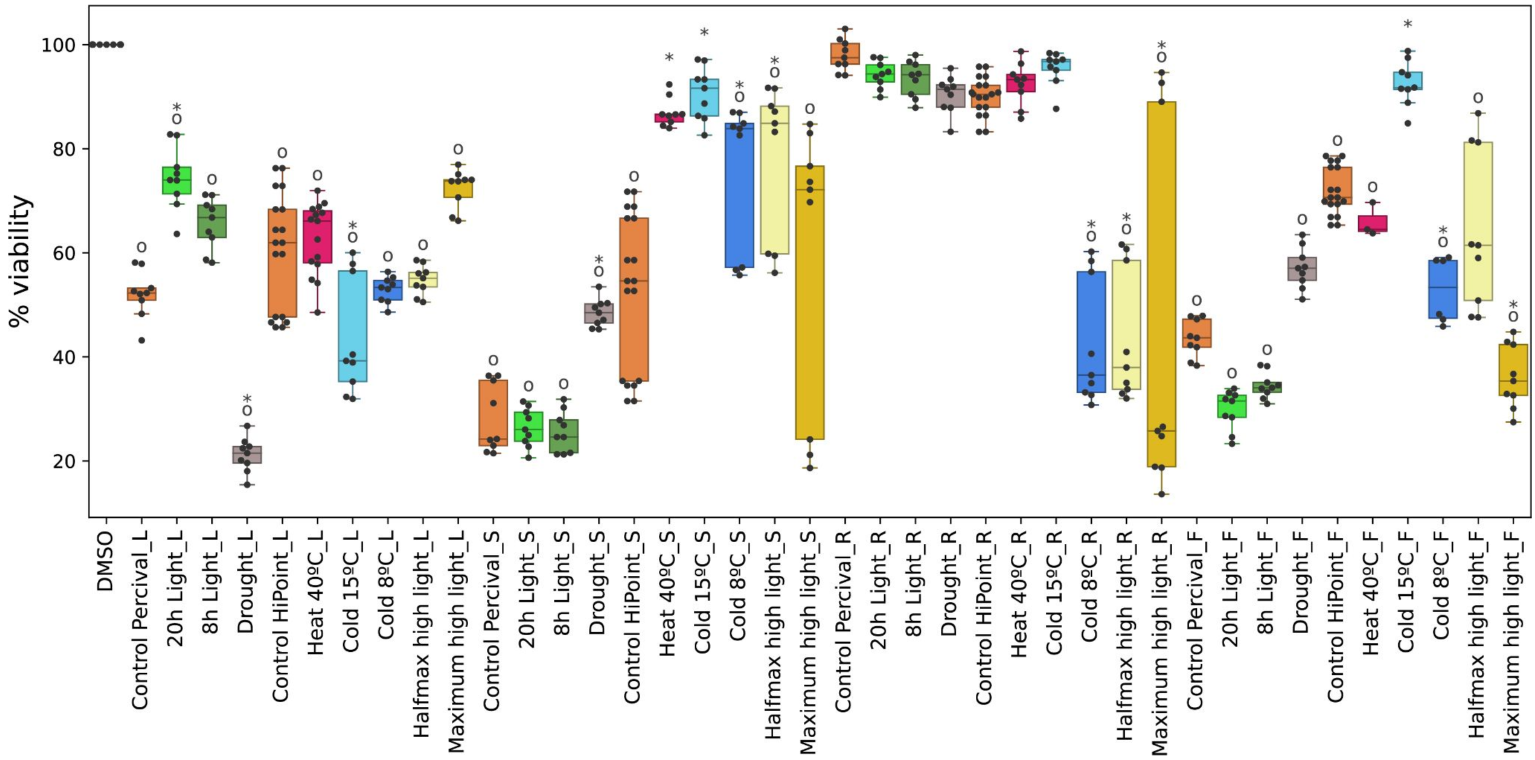


**Figure S10. MTT assay activities against SKBR3 cancer cell lines.** Samples are indicated on the x-axis, while cell viability is shown on the y-axis. The black dots indicate one measurement, while the boxplots summarize the distribution of the data. For each sample, three plants were collected, and the activity of each sampled organ was measured with three MTT assay measurements, giving nine measurements per treatment and organ. The organs are L, leaves; S, stems; F, flowers; R, roots. Circles and asterisks indicate significant differences ($p<0.05$) between the treatment and the DMSO control, and between plant extract control and treatment, respectively.

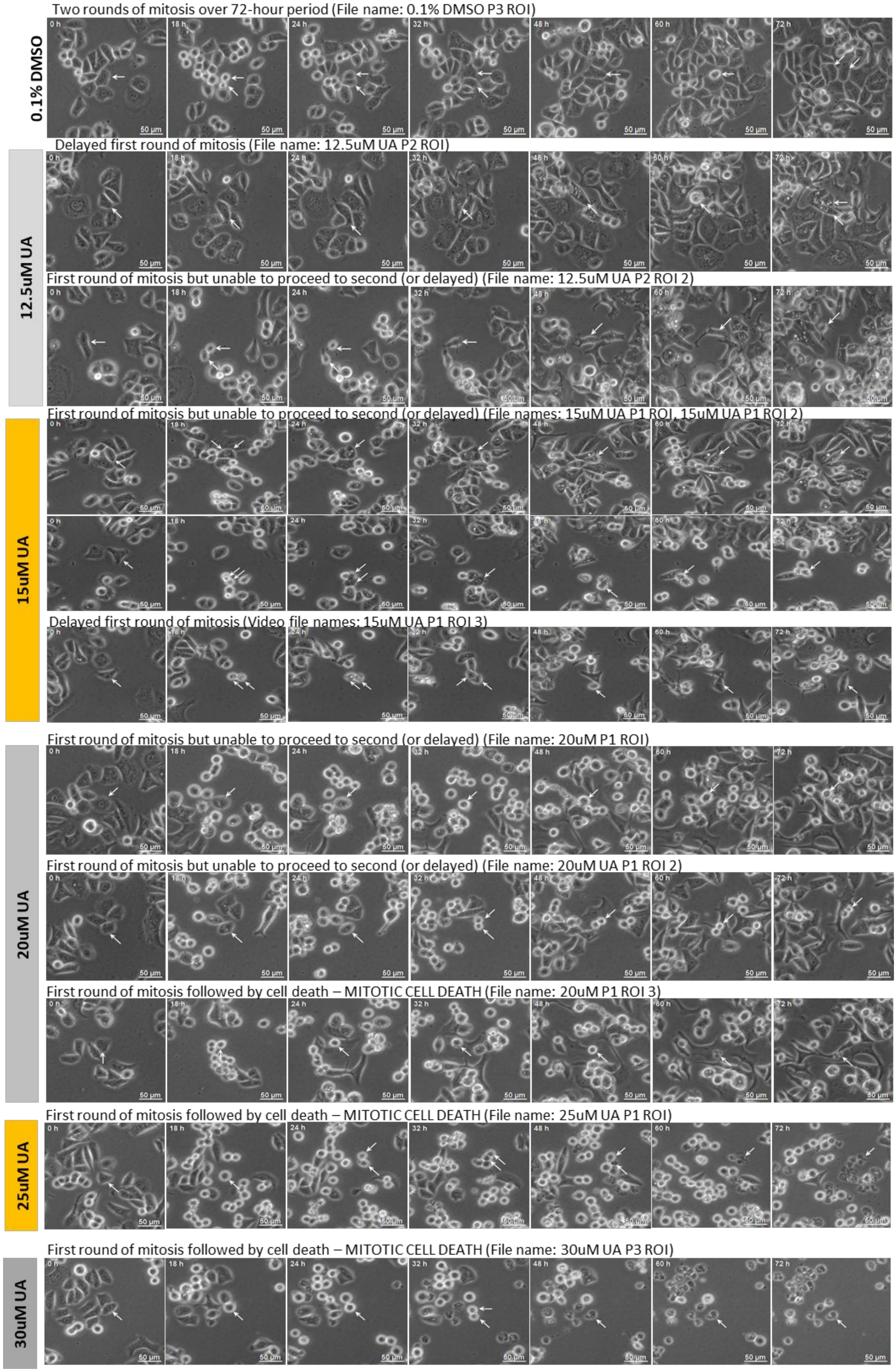


**Figure S11. Phase contrast images of SKBR3 cells treated with ursolic acid and DMSO control over 72 hours.** Rows represent different concentrations, while columns correspond to timepoints. The white arrows indicate the discussed cells.

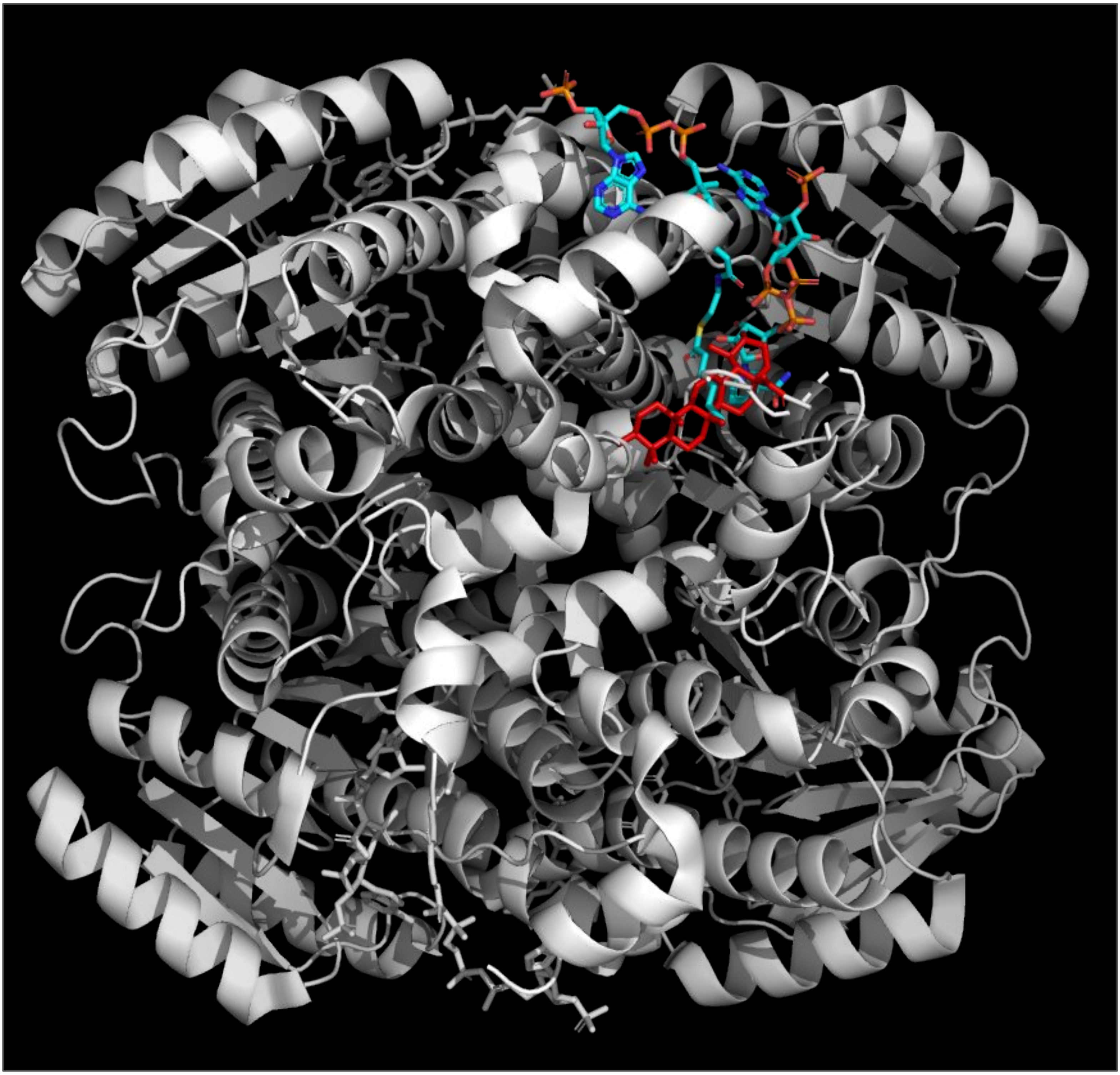

**Figure S12. Atomic model of DECR1 in complex with natural ligand and ursolic acid**. The protein is colored white, ursolic acid is colored in red, while the natural ligands are colored orange (NADP nicotiamide-adenine-dinucleotide phosphate) and cyan (hexanoyl-coenzyme A).
